# A comparative analysis of BrdU and EdU genotoxicity shows that trypanosomatids exhibit unique responses to the presence of these thymidine analogs in their DNA

**DOI:** 10.64898/2026.11.01.763445

**Authors:** Marcos Antonio F. de Oliveira, Vitor L. da Silva, Thaise L. Teixeira, Fernanda M. Gerolamo, Tarcillo J. N. Gaziri, Jennifer A. Black, Fernanda de Jesus Silva, Hernan Chaimovich, Iolanda M. Cuccovia, Marcelo S. da Silva

**Author notes:** **Corresponding author**: Marcelo S. da Silva – 748, Professor Lineu Prestes Avenue, block 9, room 916, São Paulo, SP, Brazil. Postal code: 05508-000. These authors contributed equally to this work (Marcos Antonio F. de Oliveira, Vitor L. da Silva, Thaise L. Teixeira).

## Abstract

The thymidine analogs 5-bromo-2′-deoxyuridine (BrdU) and 5-ethynyl-2′-deoxyuridine (EdU) are widely used to monitor DNA synthesis. Although studies have reported EdU toxicity in certain cell types, no reports to date describe genotoxic effects associated with BrdU/EdU use in trypanosomatids, etiological agents of neglected tropical diseases. This lack of studies raises two possibilities: either these nucleotides are not toxic to these parasites, or the observed genotoxicity was not relevant under the experimental conditions used. Here, we assessed the potential toxicity of BrdU and EdU across three pathogenic trypanosomatids: *Trypanosoma cruzi*, *Trypanosoma brucei*, and *Leishmania major.* Our findings show that EdU has greater genotoxic potential than BrdU. Nevertheless, trypanosomatids respond markedly differently to DNA alterations caused by this analog: *T. cruzi* cells stop replicating and accumulate in S phase, *T. brucei* tolerates the DNA lesions generated, while part of the *L. major* population appears to display an ‘altruistic’ behavior that benefits the survivors. Furthermore, *in silico* assays suggest that, once incorporated into DNA, the ethynyl group increases the EdU’s polarity, thereby altering DNA structure more strongly than BrdU does after the first round of DNA replication. Together, our findings suggest that using thymidine analogs in trypanosomatids should be preceded by rigorous preliminary analysis, especially when using EdU. Otherwise, the data may be biased by a BrdU/EdU-triggered DNA damage response. We believe this study may serve as an atlas to guide the initial selection of thymidine analog concentrations and exposure periods.

**Graphical Abstract:** 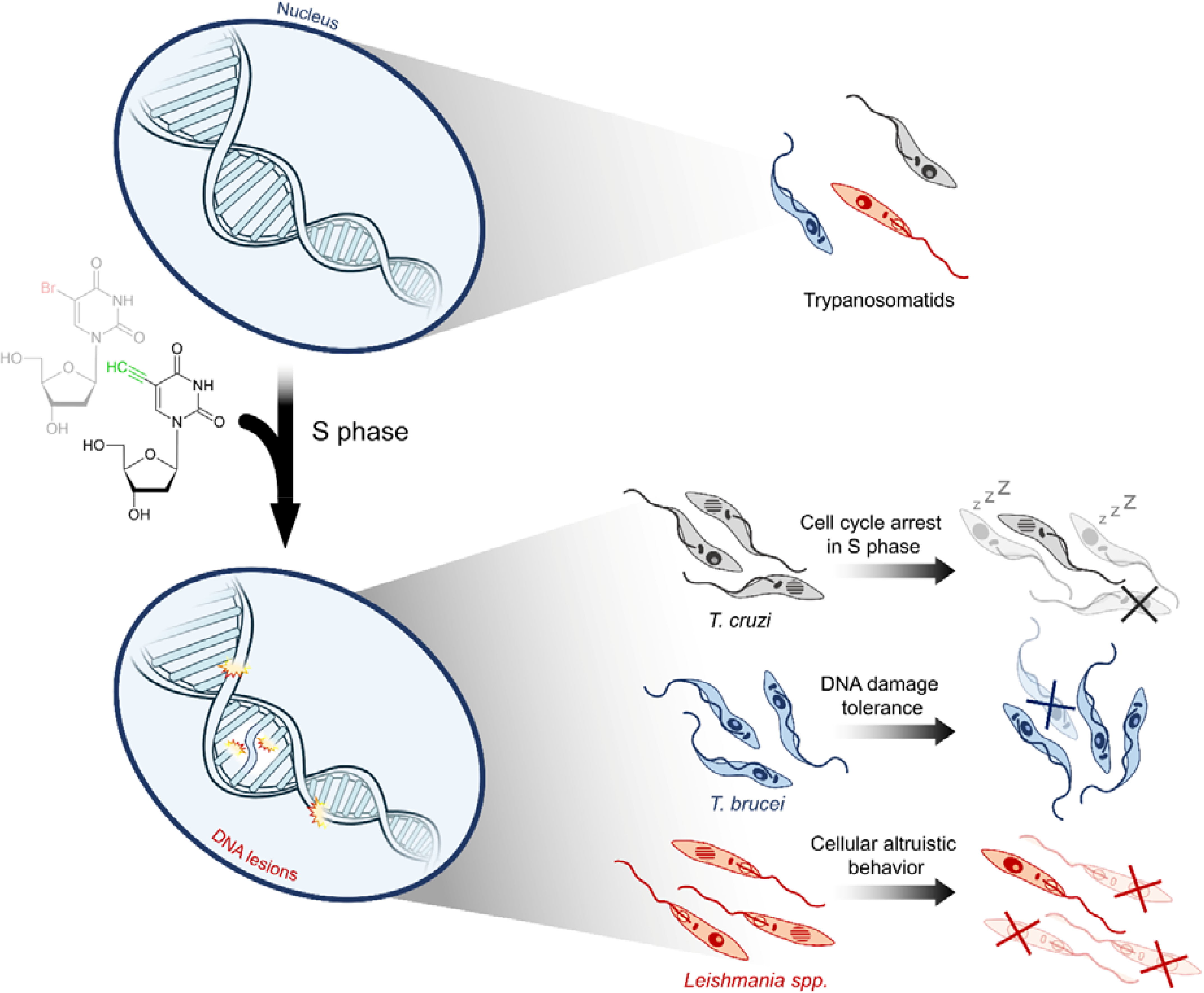

## Introduction

Trypanosomatids are single-celled eukaryotic parasites, some of which cause neglected tropical diseases (NTDs) of paramount medical relevance. These parasites include *Trypanosoma cruzi*, the etiological agent of Chagas disease (American Trypanosomiasis), *Trypanosoma brucei*, the etiological agent of Sleeping sickness (African Trypanosomiasis), and *Leishmania* spp., which cause leishmaniasis. Collectively, ∼20 million people worldwide are affected by these NTDs, amounting to ∼50,000 deaths annually^1,2,3^. Unlike model eukaryotes, some core features of trypanosomatid biology are unusually divergent, exemplified by (I) the organization of genes transcribed by RNA-pol II into large polycistronic transcription units^4^; (II) the lack of introns in most protein coding genes^5^; (III) the use of trans-splicing and polyadenylation for mRNA maturation^4^; (IV) specific thymidine modifications called bases-J, whose function is mainly associated with transcription regulation^6^, and (V) the absence of the classical DNA repair pathway non-homologous end joining (cNHEJ)^7^. Moreover, it is worth highlighting that trypanosomatids can assume different morphological forms during their life cycle in the insect and the mammalian hosts. In general, both insect and mammalian hosts harbor proliferative forms of *T. cruzi*, *T. brucei*, and *L. major*^8^. The specific terminology for these forms varies by species^8^, and the duration of the cell cycle phases can be estimated by monitoring DNA synthesis^9^.

The use of thymidine analogs is one of the primary ways to monitor DNA synthesis in different cell types^10,11^, including proliferative forms of trypanosomatids^12–14^. Over the years, the halogenated thymidine analogues, where the methyl group at position 5 is replaced by halogen atoms (F, Cl, Br or I), have become widely used, particularly BrdU (5-bromo-2’-deoxyuridine). To detect BrdU, as well as other halogenated analogs, DNA must be denatured to expose the bromo group, followed by immune detection using an α-BrdU antibody^15^. Typically, DNA denaturation is performed with harsh chemical reagents, e.g., high concentrations of HCl or NaOH, which leads to DNA instability and disruption of other cellular components.

To solve this problem and optimize the time required for detection, a new approach was developed in 2008, using Cu-catalyzed azide alkyne cycloaddition reaction (‘click’ chemistry) and a new thymidine analog called 5-ethynyl-2’-deoxyuridine (EdU)^16^. In this thymidine analog, an ethynyl group (two carbons linked by an alkyne bond) replaces the methyl group at position 5 of the base of conventional thymidine. For the detection of EdU, the click chemistry reaction is used to form a covalent bond between the ethynyl group and a fluorescent (or biotin)-labeled azide. The great advantage of using EdU is that the detection reaction is fast and sensitive and does not require DNA denaturation, which allowed EdU to become popular in the last decades^17–19^.

However, despite its widespread use, studies have reported toxicity in mammalian and yeast cells^20–22^. Although the precise molecular mechanism of EdU toxicity is yet to be established, it has been proposed that the ethynyl group may induce the formation of interstrand cross-links^21,22^, triggering the activation of the transcription-coupled nucleotide excision repair (TC-NER) pathway to remove EdU^23^. Although these and other studies have reported EdU toxicity in mammalian and yeast cells^20–24^, to date there are no reports of any toxic effects associated with the use of thymidine analogs in trypanosomatids. This lack of studies leads us to consider whether these nucleotides are non-toxic to these parasites, or whether the observed genotoxicity was not relevant under the experimental conditions used.

Thus, we sought to further evaluate the toxici effects of BrdU and EdU in three pathogenic trypanosomatids: *T. cruzi*, *T. brucei*, and *L. major*. Our findings show that EdU has greater genotoxic potential than BrdU at high concentrations (50 and 100 μM). However, the three trypanosomatids analyzed responded markedly differently to the presence of these analogs. *T. cruzi* exhibits signs of enhanced sensitivity relative to the other two parasites. This is characterized by the absence of significant proliferation following EdU exposure, coinciding with cells accumulation in S phase and a marked increase in γH2A-positive cells. *T. brucei* appears to tolerate the DNA alterations caused by the presence of these analogs at all concentrations tested, whereas *L. major* seems to possess an efficient ‘mechanism of cellular altruism’ in which, following exposure to 50 and 100 μM EdU, a portion of the population dies while the remainder resumes proliferation/viability.

To investigate the mechanism of the genotoxicity generated by these analogs, we performed *in silico* analyses. Our findings suggest that, when incorporated in DNA, EdU is more strongly associated with increases in polarity coinciding with the location of the ethynyl group than BrdU is with its corresponding bromo group. Furthermore, our molecular dynamics findings suggest that after the first round of DNA replication, EdU incorporation alters the number of hydrogen bonds (H-bonds) between DNA bases more than BrdU, suggesting negative consequences for DNA double-helix structures and stability.

## Results

### EdU impairs trypanosomatid proliferation and viability

To investigate and compare the consequences of incorporating BrdU/EdU into *T. cruzi*, *T. brucei*, and *L. major*, we first sought to assess proliferation and viability. To this end, the parasites were continuously exposed to 10 µM, 50 µM, or 100 µM of BrdU/EdU for 1 h (day 1) and 24 h (day 2). Subsequently, cells were washed and maintained in a BrdU/EdU-free culture medium until 72 h (day 5) (recovery period). Proliferation and viability were assessed immediately following the 1 h and 24 h BrdU/EdU uptake periods (exposure), as well as at 24, 48, and 72 h post-exposure (recovery period).

In general, exposure to BrdU (10, 50, or 100 µM) did not significantly alter the proliferation of the three parasites analyzed (Fig. 1A, C and E) during the periods examined. However, we observed slight decreases in viability for *T. cruzi* (after 48 h of BrdU recovery – day 4), for *T. brucei* (after 72 h of BrdU recovery – day 5), and for *L. major* (after 24 h of BrdU recovery – day 3) (Fig. 1B, D and F – right graphs).

**Fig. 1.**
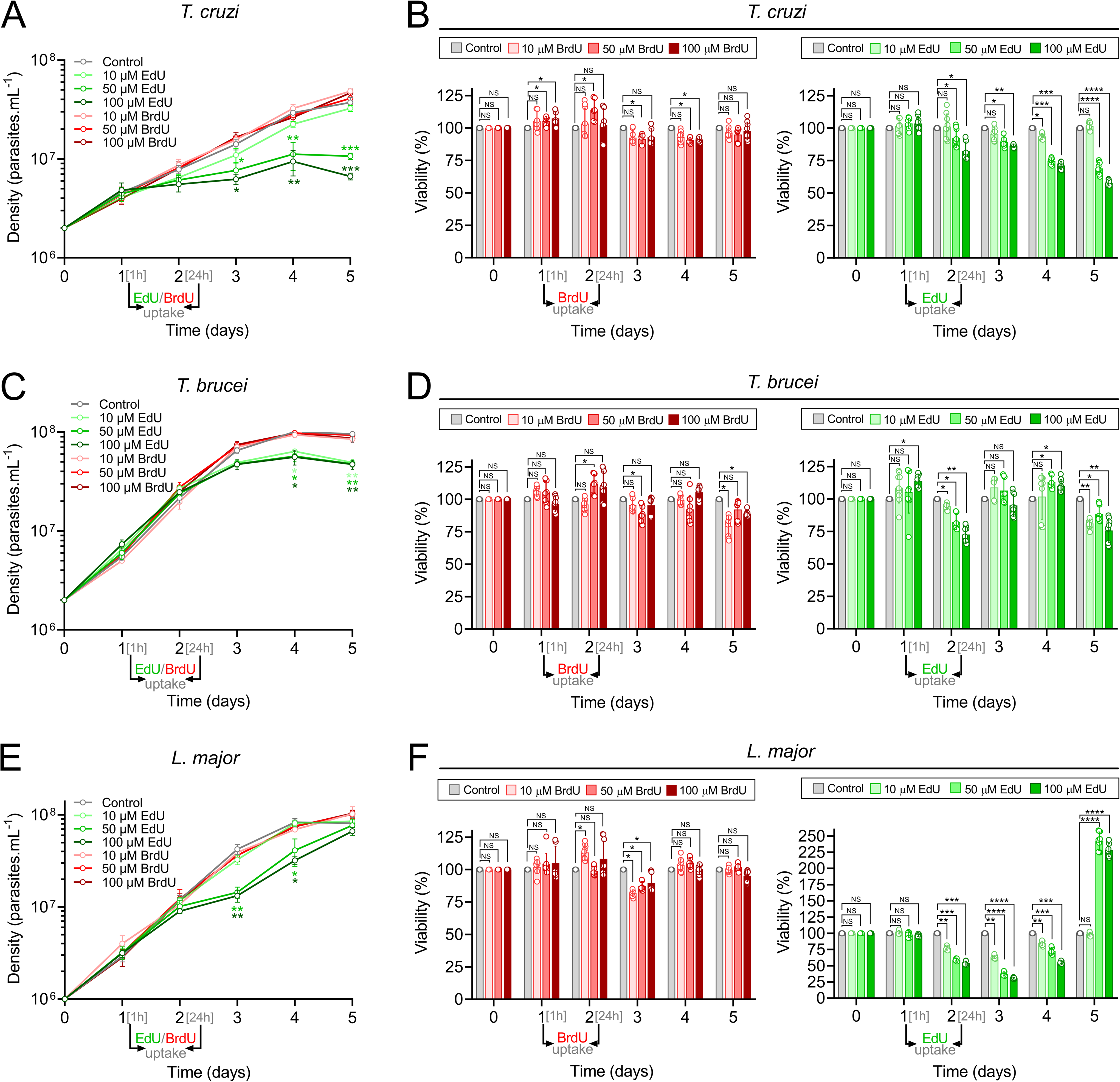
Growth curves and viability assays of trypanosomatids in response to BrdU/EdU exposure and recovery. **A**. **C** and **E**. Growth curves in the presence of different concentrations of BrdU/EdU for *T. cruzi*, *T. brucei*, and *L. major*, respectively. **B**, **D**, and **F**. Viability assay in the presence of different concentrations of BrdU (red) or EdU (green) for *T. cruzi*, *T. brucei*, and *L. major*, respectively. Assays were performed in biological triplicate. Error bars indicate the standard error of the mean (SEM). Statistical significance was assessed using Ordinary one-way ANOVA and Dunnett’s multiple comparisons. Significance is indicated as: \**p* < 0.05, \*\**p* < 0.01, \*\*\**p* < 0.001. NS denotes no significant difference according to Student’s *t*-test.

Conversely, although we did not observe a significant impact of EdU on proliferation at 10 μM in any of the parasites analyzed, EdU concentrations of 50 μM and 100 μM significantly reduced the proliferation of *T. cruzi* (starting from 24 h of recovery after EdU exposure – days 3, 4 and 5), *T. brucei* (starting from 48 h of recovery after EdU exposure – days 4 and 5), and *L. major* (between 24 and 48 h of recovery after EdU exposure – days 3 and 4) (Fig. 1A, C and E). Curiously, unlike *T. cruzi* and *T. brucei,* the *L. major* population resumed growth after 72 h of recovery (day 5) following exposure to 50 μM and 100 μM EdU, reaching the density of the untreated control (Fig.1E).

These proliferation-related phenotypic characteristics were reflected in viability assays. Groups exposed to 50 and 100 µM EdU showed a significant reduction in viability at specific time points: for *T. cruzi*, we observed a reduction in viability 24 h after EdU exposure (day 2) and during the subsequent recovery periods (days 3, 4, and 5); for *T. brucei*, we observed a slight reduction in viability right after the 24 h EdU exposure (day 2) and after 72 h of recovery (day 5); for *L. major*, we observed reduced viability 24 h after EdU exposure (day 2) and after 24 and 48 h of recovery (days 3 and 4) (Fig. 1B, D, and E). Interestingly, after 72 h of recovery from EdU exposure (day 5), the *L. major* groups treated with 50 and 100 µM EdU showed a substantial increase in viability values, exceeding the reference value of the untreated control (Fig. 1F – right graph), which corroborates the resumption of proliferation previously observed (Fig. 1E). This higher viability in *L. major* can be explained by a greater proportion of cells with replicative capacity during this period compared to the untreated control group.

Together, our findings suggest that BrdU concentrations up to 100 µM (for incorporation periods of up to 24 h), do not significantly affect the proliferation and viability of *T. cruzi*, *T. brucei* and *L. major*. On the other hand, concentrations equal to or higher than 50 µM EdU (for an incorporation period from 1 to 24 h) affect *T. cruzi*, *T. brucei*, and *L. major* proliferation and viability: EdU incorporation induces an early growth arrest in *T. cruzi* (starting right after EdU removal – day 2) and a late growth arrest in *T. brucei* (starting after 24 h of recovery – day 3). This behavior was not observed in *L. major*, which could resume proliferation after 72 h of recovery (day 5).

### EdU leads to the accumulation of T. cruzi cells in the S phase after 72 h of EdU recovery

To determine whether the arrest in proliferation and viability alterations observed coincide with defective cell cycle progression, we analyzed DNA content using flow cytometry. For this, we followed the same experimental design established previously: cells were cultured in the absence or presence of BrdU/EdU for 1 h (day 1) and 24 h (day 2), then washed and maintained in a BrdU/EdU-free culture medium, with samples being collected after 72 h of recovery from BrdU/EdU exposure (day 5).

Consistent with our proliferation and viability analyses, BrdU exposure, under all conditions tested, did not significantly alter the cell cycle phases distribution of the three trypanosomatids analyzed (Fig. 2 – left graphs; Supplementary Tables 1A-C, 2A-C, 3A-C), except for a slight increase in sub-G1 in *T. brucei* after 72 h of recovery (day 5) (Fig. 2C, D – left graphs; Supplementary Table 2C). On the other hand, EdU exposure altered the distribution of cells across cell cycle phases in a distinct manner for each parasite analyzed.

**Fig. 2.**
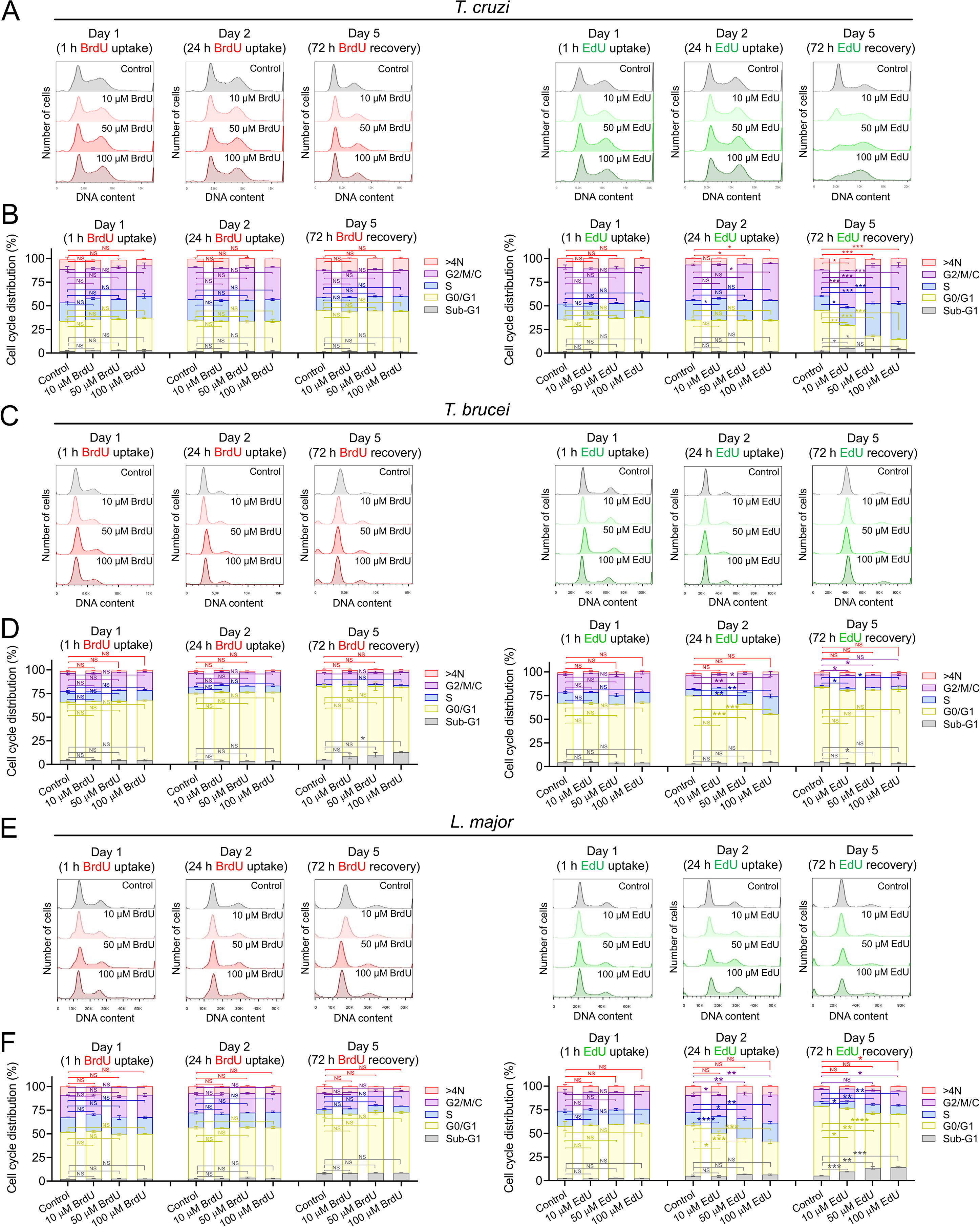
Trypanosomatid DNA content profiles in response to BrdU/EdU exposure and subsequent recovery. **A**, **C**, and **E**. Histograms from *T. cruzi*, *T. brucei* and *L. major*, respectively. The different BrdU-treated groups are represented in red. The different EdU-treated groups are shown in green. The controls are shown in gray for all groups. Assays were performed in biological triplicate. We analyzed 100,000 events per replicate. Histograms were generated using FlowJo X 10.0 software. **B**, **D**, and **F**. Quantification of the histograms shown in A, C, and E, respectively, and inference of cell cycle phases. The Y-axis represents the percentage of cells for each cycle phase (G1/G0, S, and G2/M/C), while the X-axis represents the experimental BrdU/EdU-treated groups. We performed the quantification using the “FlowJo X 10.0” software. The values and the standard error of the mean (SEM) were obtained from assays performed in triplicate. Statistical significance was assessed using unpaired two-sided Student’s t tests. Significance is indicated as: \**p* < 0.05, \*\**p* < 0.01, \*\*\**p* < 0.001. NS denotes no significant difference according to Student’s *t*-test.

For *T. cruzi*, no significant impact upon the cell cycle distribution was observed within 24 h of EdU exposure for either group (Supplementary Tables 1D, E). However, in keeping with our proliferation analysis, we observed a significant accumulation of cells in S phase at the end of a 72 h of recovery period following EdU exposure (day 5), a more modest accumulation of cells in G2/M phase, and a corresponding reduction in the proportion of G1 cells (Supplementary Table 1F; Fig. 2A, B – right graphs). Overall, the distribution of *T. cruzi* cells across cell cycle phases suggests that this parasite cannot efficiently cope with the potential alterations caused by EdU during the analyzed period (up to 72 h of recovery following EdU exposure).

Exposure to EdU (50 and 100 μM) also altered the distribution profile of *T. brucei* cells across the cell cycle phases, albeit in a more subtle way (Supplementary Tables 2D-F; Fig. 2C, D – right graphs). An examination of the cell cycle revealed only modest, but significant increases 24 h after EdU exposure (day 2) in S and G2/M phases, and an expected reduction in G1 phase (Supplementary Table 2E; Fig. 2C, D – right graphs). After 72 h of recovery period following EdU exposure (day 5), we detected no significant differences (Supplementary Table 2F; Fig. 2C, D – right graphs), suggesting that *T. brucei* can either rapidly resolve potential replication stresses or tolerate DNA damage induced by EdU.

Upon exposure of *L. major* to EdU, no significant impact upon the cell cycle distribution was observed within 1 h of EdU exposure for either group (Supplementary Tables 3D). However, a progressive accumulation of cells was detected after 24 h of EdU exposure (day 2) in S and G2/M phases, followed by an expected reduction in G1 phase (Supplementary Table 3E; Fig. 2E, F – right graphs). This cell cycle phase distribution profile was virtually maintained after 72 h of recovery period following EdU exposure (day 5), except for an increase observed in the sub-G1 population (Supplementary Table 3F; Fig. 2E, F – right graphs). This result corroborates previous findings (Fig. 1E, F) and suggests that *L. major* is more sensitive to the presence of EdU than *Trypanosoma* spp., such that a portion of its population dies after EdU incorporation (evidenced by the increased sub-G1 population), while the remainder survives and resumes proliferation.

Together, these findings suggest that 72 h is not enough time for *T. cruzi* to cope with the genotoxic alterations induced by 24 h of EdU exposure. *T. brucei* and *L. major*, on the other hand, seem to cope well (albeit in distinct ways) with the alterations caused by the presence of EdU within the analyzed timeframe.

### EdU persists in the trypanosomatid genomes after the recovery period

Given the above, we sought to investigate if the genotoxic effects of BrdU/EdU previously observed may arise due to the persistence of incorporated BrdU/EdU in the genomes of these organisms. To this end, following the same experimental design described for flow cytometry analysis, we used fluorescence microscopy to test for cells positive for nuclear BrdU/EdU after 1 h exposure (day 1), 24 h exposure (day 2), and 72 h of recovery from BrdU/EdU exposure (day 5).

For *T. cruzi*, we observed that after 1 h (day 1), BrdU is not as efficiently incorporated in DNA as EdU (Supplementary Tables 4A, D). This difference in incorporation efficiency between BrdU and EdU becomes more evident on day 2, after 24 h of exposure (Supplementary Tables 4B, E; Fig. 3A, B). However, on day 5 (after 72 h of recovery), the proportion of BrdU-positive cells decreases compared to day 2 (Supplementary Tables 4B, C), while the proportion of EdU-positive cells remains practically unchanged (Supplementary Tables 4E, F). This finding indicates that, unlike BrdU, EdU persists in the *T. cruzi* genome (Fig. 3A, B).

**Fig. 3.**
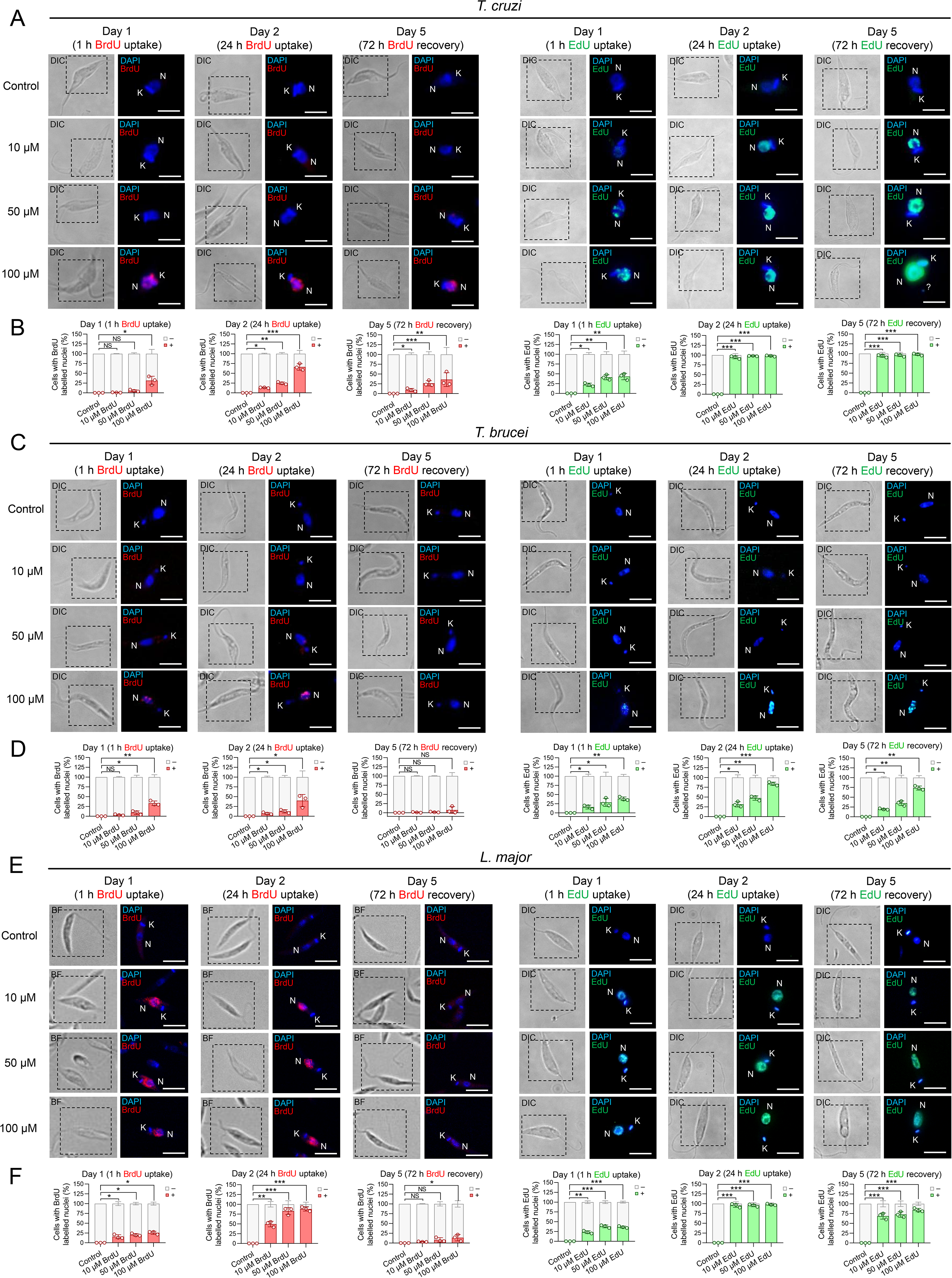
BrdU/EdU retention time in the three trypanosomatids analysed. **A**, **C**, and **E**. Representative images of the detection of BrdU (red) and EdU (green) from *T. cruzi*, *T. brucei*, and *L. major*, respectively. The bars represent 5 μm. DNA-containing organelles (nucleus – N and kinetoplast – K) were visualized using DAPI (blue). DIC stands for differential interference contrast. The quantification of the proportion of BrdU/EdU-positive cells was expressed as a percentage of the total number of cells counted and is shown in **B**, for *T. cruzi*, **D**, for *T. brucei,* and **F**, for *L. major*. The values and the standard error of the mean (SEM) were obtained from assays performed in triplicate. Statistical significance was assessed using unpaired two-sided Student’s t tests. Significance is indicated as: \**p* < 0.05, \*\**p* < 0.01, \*\*\**p* < 0.001. NS denotes no significant difference according to Student’s *t*-test.

In *T. brucei*, we observed a behavior similar to *T. cruzi* after 1 h (day 1) and 24 h of exposure (day 2), i.e.: BrdU is not as efficiently incorporated in DNA as EdU (Supplementary Tables 5A, B, D, E; Fig. 3C, D). Interestingly, after 72 h of recovery, the proportion of BrdU-positive cells significantly decreases (Supplementary Table 5C) compared to day 2 (Supplementary Table 5B), while in the same period, the proportion of EdU-positive cells only slightly decreases (Supplementary Table 5F) compared to day 2 (Supplementary Table 5E; Fig. 3C, D). This finding suggests that EdU also persists in the *T. brucei* genome, though not to the same proportion as observed in *T. cruzi*.

For *L. major*, we observed a behavior similar to that of found in *T. brucei* and *T. cruzi* (Supplementary Tables 6A-F; Fig 3E, F). However, compared with *Trypanosoma* spp. (Supplementary Tables 4B, 5B), *L. major* showed a higher proportion of BrdU-positive cells after 24 h of exposure (Supplementary Table 6B), suggesting greater BrdU incorporation. Nevertheless, even with this high proportion of BrdU-positive cells, BrdU is still not incorporated in DNA as efficiently as EdU after 24 h of exposure (Supplementary Tables 6B, E). After 72 h of recovery (day 5), the proportion of BrdU-positive cells decreases considerably (Supplementary Table 6C) compared to day 2 (Supplementary Table 6B), reaching levels similar to those previously observed for *T. brucei* (Supplementary Tables 5C). In the same period (after 72 h of recovery), the proportion of EdU-positive cells slightly decreases (Supplementary Table 6F) compared to day 2 (Supplementary Table 6E; Fig. 3E, F). This finding suggests that EdU also persists in the *L. major* genome at a relative proportion similar to that observed in *T. brucei* (Supplementary Table 5F).

Across all three trypanosomatids analyzed, we observed that BrdU is incorporated/detected in DNA less efficiently than EdU. In contrast, its removal from the genome appears to be more efficient than EdU’s. It is worth noting that the decrease in the percentage of EdU-positive cells between day 2 and day 5 for *T. brucei* (↓38.1%, ↓26.9%, and ↓12.5%, respectively for 10, 50 and 100 μM EdU) and *L. major* (↓27.9%, ↓23.78%, and ↓14%, respectively for 10, 50 and 100 μM EdU) was much greater relative to *T. cruzi* (↓0.35%, ↓2.22%, and ↓0%, respectively for 10, 50 and 100 μM EdU) (Supplementary Tables 4E, 4F, 5E, 5F, 6E, 6F). Given that the recovery period was 72 h post-EdU exposure and the doubling time of *T. cruzi* is ∼24 h^9^, this EdU persistence can be explained if *T. cruzi* cells are not committed to the cell cycle during the recovery period, which is corroborated by the decrease in proliferative rate observed in the growth curves (Fig. 1A) and the cell cycle alterations observed in flow cytometry analysis (Fig. 2A, B). However, we cannot ignore the fact that the doubling time of *T. brucei* and *L. major* is approximately 3 times shorter (∼8 h)^9,25^ than that of *T. cruzi*^9^, which could account for the persistence of EdU in a larger proportion of *T. cruzi* cells compared to the other trypanosomatids.

The differences observed in the incorporation and persistence/clearance of BrdU and EdU within the genomes of these trypanosomatids (across the exposure and recovery periods) raise the question of whether the presence of these analogs could cause DNA damage in these organisms.

### T. cruzi and L. major exhibit proportionally increased DNA damage signaling in response to the different EdU concentrations

To assess whether BrdU/EdU incorporation coincides with increased DNA lesions in these three trypanosomatids, we examined histone H2A phosphorylation^26^, using the same experimental design described earlier for flow cytometry and fluorescent microscopy. It is worth mentioning that phosphorylated histone H2A (γH2A) is one of the earliest markers of DNA damage in trypanosomatids and its detection via immunofluorescence assay (IFA) using an α-γH2A antiserum has been commonly employed in *T. cruzi*^19^, *T. brucei*^26^, and *L. major*^27^.

For *T. cruzi*, we did not observe significant alterations in the percentage of γH2A-stained (positive) nuclei across most BrdU concentrations during the exposure period (days 1 and 2) (Supplementary Tables 7A, B). However, after 72 h of recovery from BrdU exposure, we observed a slightly increased proportion of γH2A-labeled cells, followed by slightly increased γH2A intensity (Supplementary Table 7C; Fig. 4A-C – left graphs). For the EdU-treated groups, we observed no significant differences in the proportion of γH2A-labeled cells after 1 h EdU uptake, and slight increases in γH2A intensity proportional to increasing EdU concentrations (Supplementary Table 7D; Fig. 4A-C – right graphs). This finding indicates that exposure to 10 – 100 µM EdU for 1 h does not alter the number of cells signaling DNA damage, but it modestly and proportionally increases the number of cells with higher γH2A signal intensity, reflecting more DNA lesions proportional to EdU concentration. Nevertheless, at 24 h of EdU exposure (day 2) and 72 h of recovery from EdU exposure (day 5), we noted a significant increase in both the number of γH2A-labeled cells, and the overall nuclear intensity of γH2A (Supplementary Tables 7E, F; Fig. 4A-C – right graphs). These alterations were proportional to the EdU concentrations used, i.e., the higher the concentration of EdU, the greater the percentage and intensity of γH2A-labeled cells (Fig. 4A-C – right graphs). These findings align with what was previously suggested and indicate that the persistence of EdU in the *T. cruzi* genome is being recognized as a DNA lesion.

**Fig. 4.**
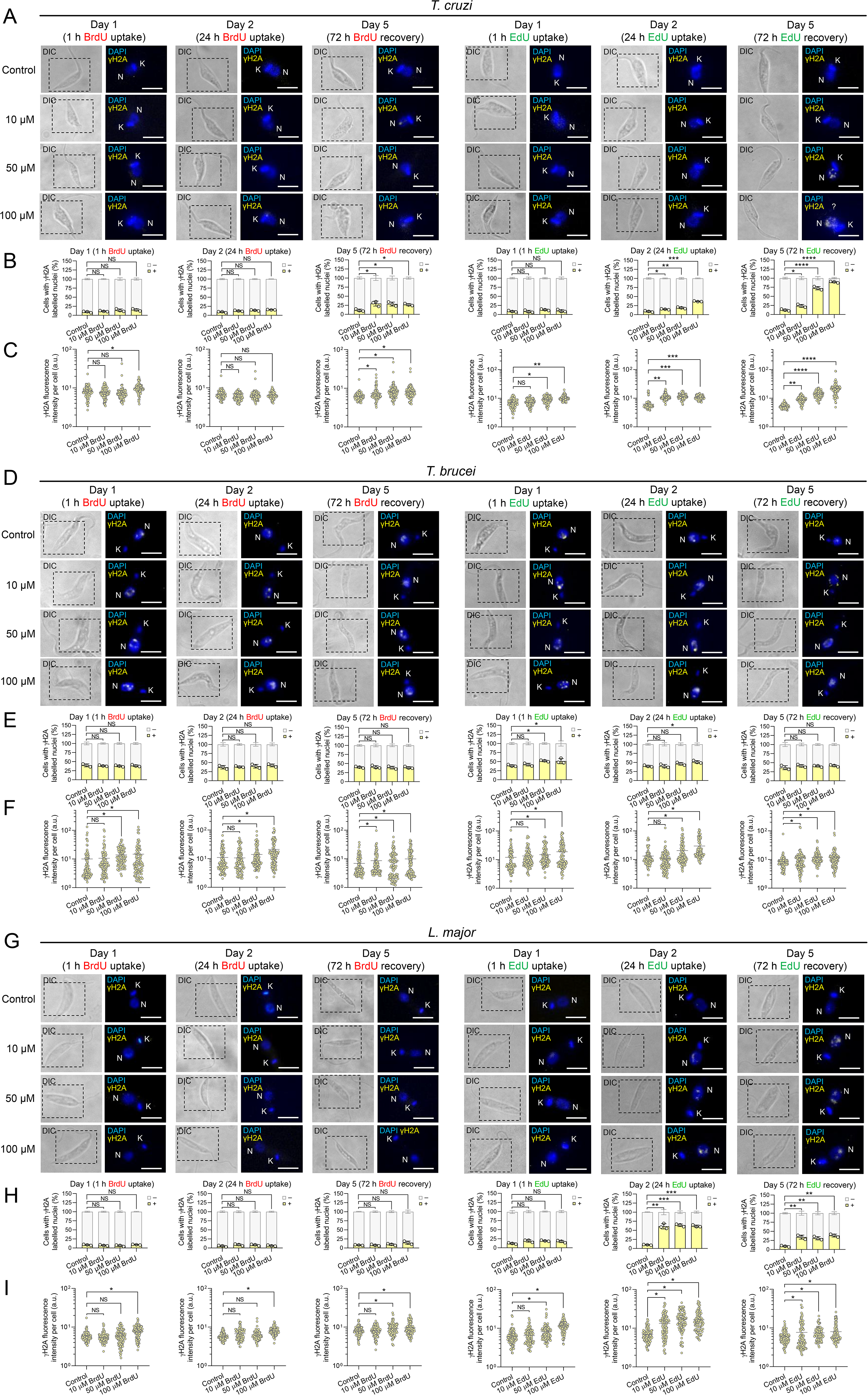
DNA damage signaling detection using γH2A. **A**, **D**, and **G**. Representative images of the fluorescence signal for γH2A (yellow) from *T. cruzi*, *T. brucei*, and *L. major*, respectively. DAPI (blue) stains the DNA-containing organelles (nucleus – N and kinetoplast – K). DIC stands for differential interference contrast. The bar represents 5 μm. The quantification of the proportion of γH2A-positive cells (%) is shown in **B**, for *T. cruzi*, **E**, for *T. brucei,* and **H**, for *L. major*. The quantification of γH2A fluorescence intensity in the nuclei (in arbitrary units – a.u.) is shown in **C**, for *T. cruzi*, **F**, for *T. brucei,* and **I**, for *L. major*. The values and the standard error of the mean (SEM) were obtained from assays performed in triplicate. Statistical significance was assessed using unpaired two-sided Student’s t tests. Significance is indicated as: \**p* < 0.05, \*\**p* < 0.01, \*\*\**p* < 0.001. NS denotes no significant difference according to Student’s *t*-test.

Furthermore, it is worth highlighting that after 72 h of recovery, we observed a high proportion of *T. cruzi* cells presenting a DNA-containing micronucleus-like structure in all EdU-exposed groups relative to BrdU-exposed groups (Supplementary Fig. 1). Micronucleus emergence has been described as a critical, downstream consequence of unrepaired or misrepaired DNA Interstrand Crosslinks (ICLs)^28,29^, a type of DNA lesion resulting from the presence of EdU^20,21^. However, our study cannot confirm whether this type of DNA lesion occurs in *T. cruzi* following EdU exposure.

For *T. brucei*, we also did not observe significant changes in the percentage of γH2A-positive nuclei at any of the BrdU/EdU concentrations, across all time points analyzed (days 1, 2, and 5) (Supplementary Tables 8A-C). However, we observed a slight increase in γH2A intensity in groups exposed to 50 and 100 µM BrdU compared with the control on day 1 (Supplementary Table 8A), on day 2 (Supplementary Table 8B), and on day 5 (Supplementary Table 8C) (Fig. 4D-F – left graphs). A similar profile of increased γH2A intensity in groups exposed to 50 and 100 µM EdU was also observed on day 1 (Supplementary Table 8D), on day 2 (Supplementary Table 8E), and on day 5 (Supplementary Table 8F) (Fig. 4D-F – right graphs). These modest alterations suggest that *T. brucei* exhibits DNA damage tolerance in response to the presence of BrdU/EdU in its genome, whether due to an absence of DNA lesions caused by BrdU/EdU or a lack of signaling in response to them. Nevertheless, an observation worth highlighting is the high proportion of cells with γH2A-stained (positive) nuclei in the control (non-exposed) groups (Supplementary Table 8; Fig. 4D-F). This finding suggests that the *T. brucei* cells used in our analysis exhibit an elevated basal level of signaling related to endogenous DNA damage.

For *L. major*, we notice for BrdU-exposed groups a profile similar to that of *T. brucei*: no significant changes in the percentage of γH2A-positive nuclei at any of the BrdU concentrations, across all time points analyzed (days 1, 2, and 5), and a slight increase in γH2A intensity in groups exposed to 100 µM BrdU compared with the control on day 1 (Supplementary Table 9A), on day 2 (Supplementary Table 9B), and on day 5 (Supplementary Table 9C) (Fig. 4G-I – left graphs).

For *L. major* EdU-treated groups, we observed no significant differences in the proportion of γH2A-labeled cells after 1 h EdU uptake, and slight increases in γH2A intensity in groups exposed to 50 and 100 µM EdU (Supplementary Table 9D) (Fig. 4I – right graphs). As in the case of *T. cruzi*, this finding indicates that exposure to 10 – 100 µM EdU for 1 h does not alter the number of cells exhibiting DNA damage signaling (γH2A), but it does, modestly and in proportion to the EdU concentration, increase the number of cells with a higher intensity of the γH2A signal (Supplementary Table 9D). Nevertheless, at 24 h of EdU exposure (day 2) and 72 h of recovery from EdU exposure (day 5), we noted a significant increase in both the number of γH2A-labeled cells, and the overall nuclear intensity of γH2A (Supplementary Table 9E, F). These alterations are greater after 24 h of EdU exposure (day 2) for all concentrations (10 – 100 μM) (Fig. 4G-I – right graphs). In principle, the decrease in the proportion of *L. major* cells presenting γH2A-positive nuclei after 72 h of recovery from EdU exposure (day 5) could suggest that *L. major* can repair the DNA lesions caused by the presence of EdU. However, this may not be the most parsimonious hypothesis, given that it does not explain the observed alterations in viability (Fig. 1F). Thus, it is more likely that a portion of the *L. major* population died due to DNA lesions generated by EdU during the recovery period, while the surviving population resumed proliferation, which is consistent with previous observations (Figures 1E, 1F, 2E, 2F).

Altogether, our data indicate that EdU exhibits greater genotoxic potential than BrdU, particularly during prolonged exposure periods (up to 24 h) and at concentrations ranging from 50 to 100 µM. However, the three trypanosomatids analyzed respond differently to the presence of these analogs in terms of DNA damage signaling, which raises questions regarding the type of DNA lesions/alterations generated by the presence of these analogs and why EdU is more genotoxic than BrdU.

### The chemical properties of BrdU and EdU can explain differences in their reactivity

To begin addressing questions about the difference in genotoxicity between BrdU and EdU, we analyzed the chemical structures of thymidine, BrdU, and EdU in depth.

Despite their similarity, analysis of electronic charge distributions on the molecular surfaces shows that BrdU’s distribution is more similar to that of dT than to that of EdU (Fig. 5A). We observed a slight increase in the phase of the molecular orbitals (φ – red) near carbon 5 after HOMO and LUMO analysis, possibly due to the electronic polarization induced by bromo/ethynyl group. Moreover, through MEP analysis we observed the molecular shape of the bromine atom linked to carbon 5 of the base is more rounded than the shape of the methyl group in dT. Although both ligands virtually present a neutral charge distribution (greenish color), the bromine atom region shows a less intense greenish color (closer to yellow), indicating a region with negative polarity (Fig. 5A). As methyl is a nonpolar group, this slight difference can be explained due to the electronegativity of the bromine atom (2.96 using the Pauling scale)^30^.

**Fig. 5.**
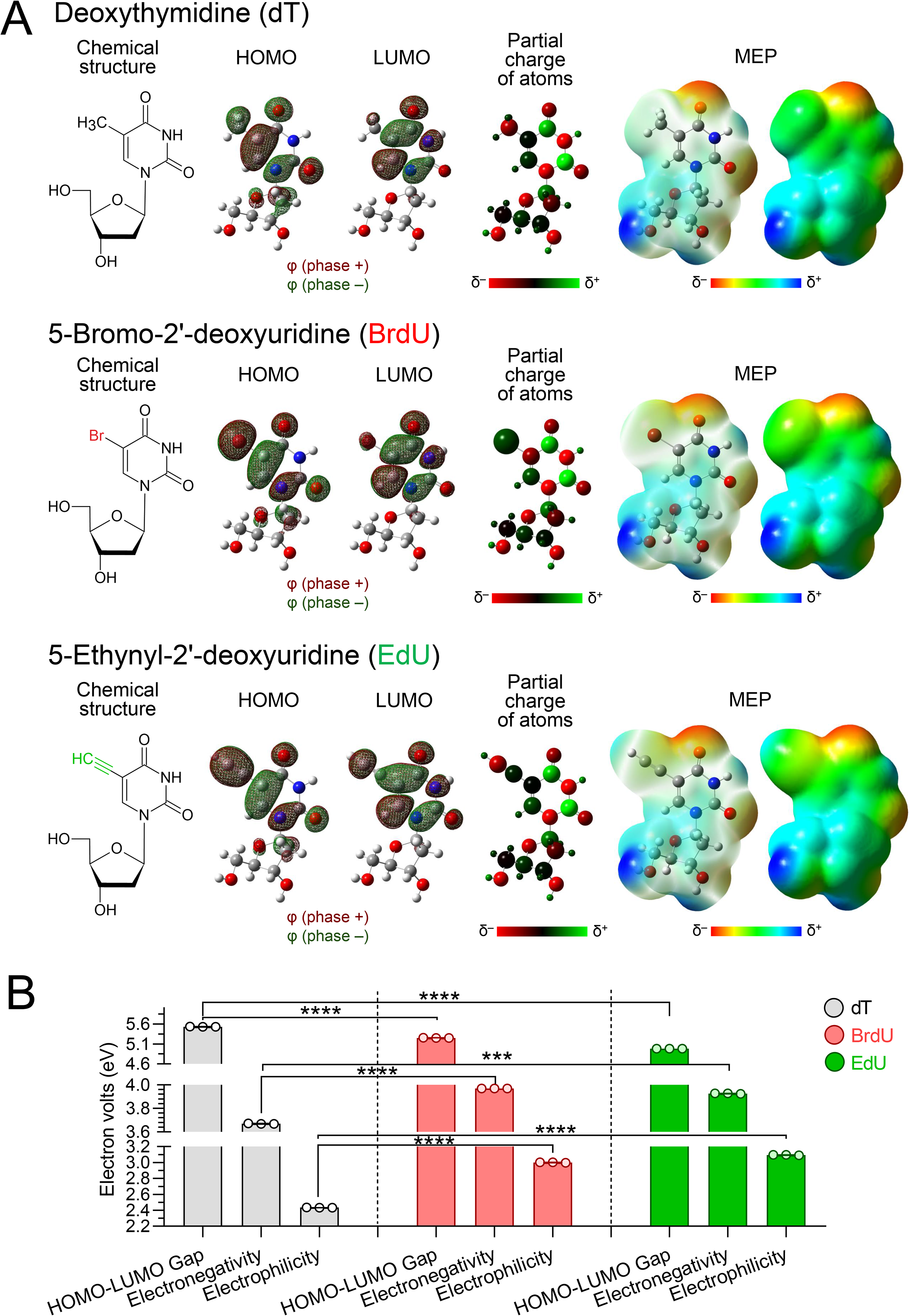
Chemical characteristics of the nucleosides dT (deoxythymidine), BrdU (5-Bromo-2’-deoxyuridine), and EdU (5-Ethynyl-2’-deoxyuridine). A. From left to right, the chemical structure, the highest occupied molecular orbital (HOMO), the lowest unoccupied molecular orbital (LUMO), the partial atomic charges, and the molecular electrostatic potential (MEP) are presented for each nucleoside analyzed (dT, BrdU, and EdU). Phi (φ) represents the phase of the molecular orbitals: red for positive and green for negative. δ is used to show partial positive (δ+) or partial negative (δ-) charges. B. Comparative analysis of the values (in mV) of HOMO-LUMO gap, electronegativity, and electrophilicity for dT, BrdU, and EdU. The error bars represent the standard error of the mean (± SEM) from an assay performed in triplicate. Statistical significance was assessed using unpaired two-sided Student’s t tests. Significance is indicated as: \**p* < 0.05, \*\**p* < 0.01, \*\*\**p* < 0.001. NS denotes no significant difference according to Student’s *t*-test.

When comparing dT to EdU, we observe that the overall electron distribution of both molecules is similar. However, when analyzing the ethynyl group linked to carbon 5, we noticed a prominent cylindrical molecular shape, which could potentially generate discrepancies in the expected number of H-bonds and consequent instability of the DNA structure. Furthermore, the presence of the triple bond between carbons in the ethynyl group (where 6 electrons are shared) presents a less intense greenish color relative to dT/BrdU, indicating high electron density in the σ-π-π bond region. Also, we observed a low electron density (δ) with a bluish color corresponding to the nitrogen bound to the carbon at the extremity of the ethynyl group (Fig. 5A).

Additionally, we compared the HOMO-LUMO gap, electronegativity, and electrophilicity values. The HOMO-LUMO gap parameter represents the energy difference between HOMO and LUMO in a molecule and may be used to determine the compound’s chemical reactivity and stability^31^. A large gap indicates higher stability and lower reactivity, whereas smaller gaps suggest that the molecule is more reactive, and its electrons are more easily polarized/excited. The HOMO-LUMO gap measured in our analysis showed decreasing values as follows: dT (5.53 eV ±0.005) > BrdU (5.25 eV ±0.005) > EdU (4.98 eV ±0.001), suggesting EdU is more reactive than BrdU and dT (Fig. 5B).

We next examined the electronegativity of each molecule, or the tendency of an atom/molecule to attract electrons to itself when participating in a chemical bond: the higher the electronegativity, the greater the tendency to attract electrons^32^. The electronegativity measured in our analysis showed values in the following order: BrdU (3.97 eV ±0.001) > EdU (3.93 eV ±0.002) > dT (3.67 eV ±0.002), indicating both BrdU and EdU have a higher tendency to attract electrons than dT (Fig. 1B), supported further by an examination of the electrophilicity of each molecule (i.e., the molecule’s affinity for electrons)^33^. This analysis supports a higher tendency of both EdU and BrdU to accept electrons when compared to dT (Fig. 1B): EdU (3.09 eV ±0.004) > BrdU (3.00 eV ±0.004) > dT (2.43 eV ±0.001).

Together, replacement of the methyl group with either a bromo (in the case of BrdU) or ethynyl (in the case of EdU) group is predicted to induce a polarity alteration. Interestingly, the polarity of the ethynyl group seems to be substantially higher relative to the bromo group, implying higher reactivity and suggesting greater genotoxicity.

### When incorporated, both BrdU and EdU alter DNA topology

To investigate further the impact of BrdU or EdU on the DNA molecule, we performed molecular dynamics simulations using hypothetical DNA molecules (20 mer) generated to emulate the incorporation of BrdU or EdU after the first (BrdU/EdU in one strand) and second (BrdU/EdU in both strands) rounds of DNA replication (Fig. 6A).

**Fig. 6.**
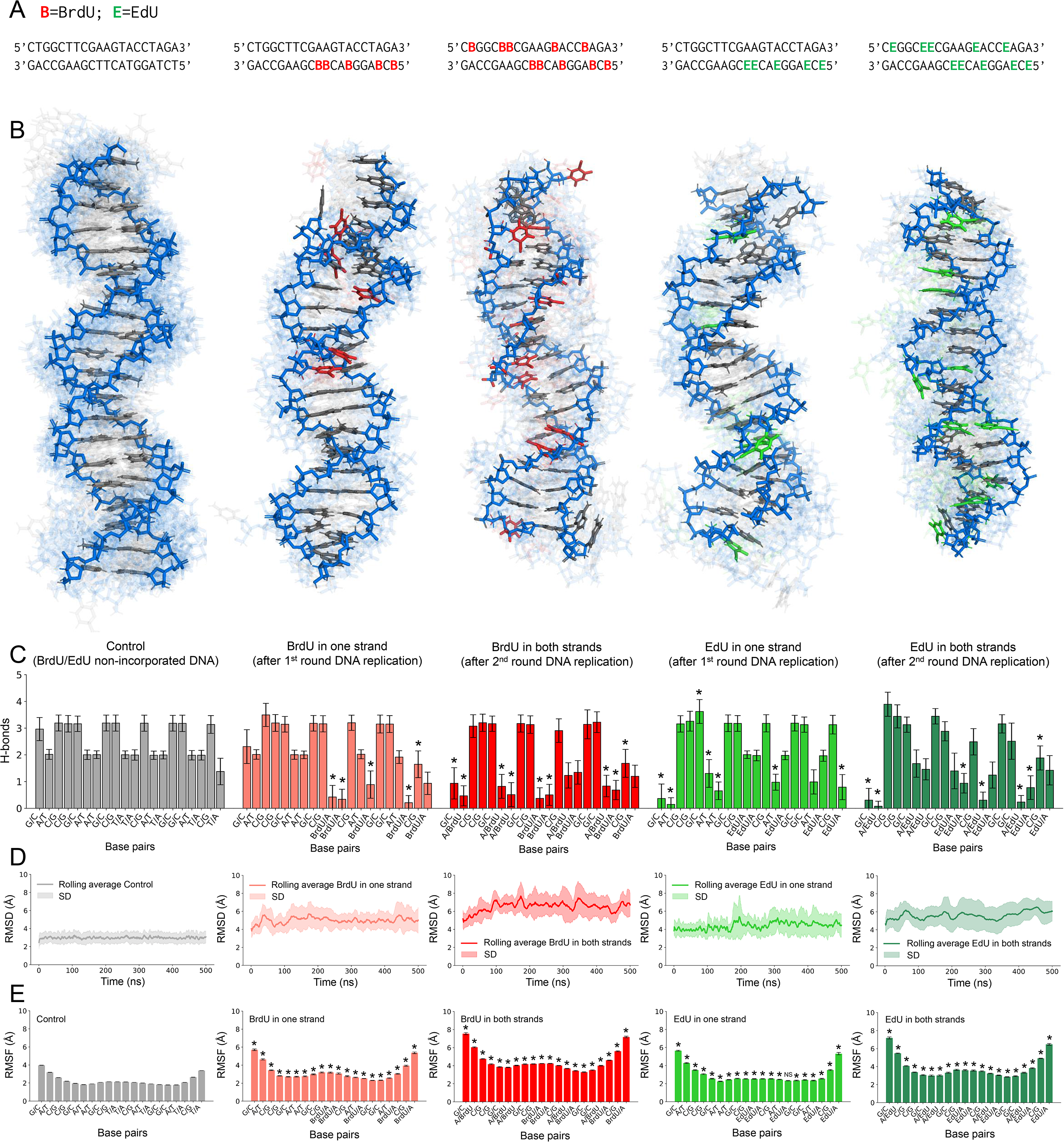
Molecular dynamics simulations reveal alterations in DNA topology in response to BrdU/EdU incorporation. **A.** DNA sequences (20 mer) used for molecular dynamics simulations. **B**. All DNA trajectories clustered into a single picture with emphasis on the most common one. From left to right: Control, BrdU incorporated in one strand, BrdU incorporated in both strands, EdU incorporated in one strand, and EdU incorporated in both strands. **C**. Bar graphs representing the average of H-bonds formed among the indicated base pairs in each simulation. The incorporation of BrdU and EdU increased RMSD (**D**) and RMSF (**E**) values. The values and the standard error of the mean (SEM) were obtained from assays performed in triplicate. Statistical significance was assessed using unpaired two-sided Student’s t tests and ordinary one-way ANOVA. Significance is indicated as: \**p* < 0.05, \*\**p* < 0.01, \*\*\**p* < 0.001. NS denotes no significant difference.

After 100 ns, all DNA trajectories were clustered into a single picture with emphasis on the most common one (Fig. 6B). The structure of the control DNA remained virtually stable during the simulation, indicating that our system was reliable and compatible with literature^34^. The simulations containing BrdU in one strand resulted in the loss of base pairing, largely coinciding with the locations of incorporated BrdU. The simulations containing BrdU in both strands resulted in unpaired BrdU nucleotides and severe double helix distortions, especially in the extremities of the DNA. For EdU, the simulations containing this analog in one strand did not result in loss of base pairing, as observed with BrdU, but enhanced double helix distortion. On the other hand, the simulations containing EdU in both strands resulted in severe double helix distortions (mainly in extremities) and unpaired EdU nucleotides, like those observed for BrdU in both strands (Fig. 6B).

Together, the incorporation of BrdU or EdU in only one DNA strand produced unpaired BrdU/EdU and fewer double helix distortions relative to the DNA containing BrdU/EdU in both strands. Moreover, our data predict that a second round of DNA replication tends to be much more damaging in the presence of BrdU/EdU.

### Both BrdU and EdU alter the number of base pairs presenting an altered number of H-bonds

In our simulations, we also evaluated the number of H-bonds formed between the bases for each situation previously described. The control DNA simulation displayed the expected number of H-bonds for each type of base pair (with A-T forming two and C-G forming three H-bonds), confirming that our system is reliable and compatible with existing data^34^.

When BrdU/EdU was incorporated into one DNA strand (1^st^ round of DNA replication), there was a significant increase in the number of base pairs presenting an altered number of H-bonds. Specifically, a greater number of base pairs underwent changes in the number of H-bonds due to the presence of EdU than due to the presence of BrdU: 7 base pairs underwent significant changes in the average number of H-bonds due to the presence of EdU, compared to only 5 due to the presence of BrdU (Fig. 6C). Interestingly, in the analysis involving BrdU, 4 of the 5 base pairs that exhibited changes in the number of H-bonds were BrdU-A pairs. In the case of EdU, only 1 of the 7 base pairs that showed changes in the number of H-bonds was an EdU-A pair. The other altered base pairs (with one exception) were not adjacent to the EdU incorporation site (Fig. 6C).

When BrdU/EdU was incorporated into both strands (2^nd^ round of DNA replication), the number of base pairs exhibiting an altered number of H-bonds increased for BrdU (from 5, when BrdU is present in only one DNA strand, to 9, when it was present in both strands), whereas it remained the same for EdU (7, whether EdU is present in only one DNA strand or in both).

In general, the number of H-bonds for each altered base pair due to the presence of BrdU/EdU was significantly lower than that observed in the control groups (Fig. 6C). Interestingly, after the 2^nd^ round of DNA replication, we noticed that some C-G base pairs in the vicinity of the BrdU/EdU-A base pairs displayed a lower number of H-bonds (Fig. 6C). We believe that this may suggest the presence of covalent bonds linking two base pairs on opposite DNA strands, a characteristic of ICLs^20,21^. However, further experiments are needed to support this hypothesis.

### BrdU and EdU yielded high RMSD and RMSF values during the simulations

From our simulations, we also obtained values for RMSD (Root Mean Square Deviation) and RMSF (Root Mean Square Fluctuation), which reflect the average distance between the backbone atoms of the superimposed DNA molecules and the average deviation of a nucleotide’s position over time, respectively (Fig. 6D, E).

As expected, control DNA (gray) showed low RMSD and RMSF values with minimal fluctuation across all nucleotides. The presence of BrdU/EdU in one strand (light red and light green, respectively) increased both the RMSD and RMSF values relative to the control. However, the presence of BrdU/EdU in both strands (dark red and dark green, respectively) increased the RMSD and RMSF values even more relative to the control and to DNAs with BrdU/EdU in one strand (Fig. 6D, E). As expected, these values were higher at the DNA extremities (Fig. 6D, E).

### DNA dihedral angles are slightly altered due to the presence of BrdU/EdU

Lastly, we examined the dihedral angles of our simulated DNA in the presence and absence of incorporated BrdU or EdU. Dihedral angles can be defined as torsional angles that describe the rotation around bonds in the DNA molecule (Supplementary Fig. 2A), playing a crucial role in its overall structure and flexibility.

When compared to the control DNA molecules, we observed the following alterations: for the α dihedral angle, we observed an increased peak at ∼60° in the simulations with BrdU/EdU in both strands, which reflects the presence of helix distortions since this angle is ∼290° in the control; for the β dihedral angle, we did not observe significant alterations; for the γ dihedral angle, we observed two small peaks at ∼180° and ∼300° in the simulations with BrdU/EdU in one and in both strands, which also reflect the presence of possible helix distortions since this angle is ∼50° in the control; for the δ dihedral angle, we noticed an increased frequency at ∼90° relative to the control, only for the simulations with BrdU (in one or in both strands); for the ε dihedral angle, we did not observe significant alterations, although there was an increased SD in the simulations with BrdU/EdU in one strand; finally, for the ζ dihedral angle, we observed an additional peak at ∼80° in the simulations with BrdU/EdU in both strands, which indicates helix distortions since this angle is absent in the control group (Supplementary Fig. 2B). It is worth highlighting that the angle frequency distribution of all dihedral angles obtained in our simulations corroborates the data obtained by other research groups^34^.

Considering the standard deviation and the new frequency generated for specific angle values, our simulations suggest that the presence of BrdU/EdU in one or both DNA strands (i.e., following the first or second round of DNA replication) causes distortions in the DNA double helix. Interestingly, this is also a characteristic intrinsically associated with the presence of ICLs^35,36^. However, further assays are needed to investigate whether ICLs are generated by the presence of BrdU/EdU in the genome of trypanosomatids.

## Discussion

BrdU and EdU have been routinely used to monitor DNA synthesis across a wide range of cell types. The concentration values used and the duration of incorporation vary depending on the intrinsic cellular characteristics of the organism in which DNA synthesis is being monitored. Some research groups observed, in mammalian cell lines, distinct incorporation efficiencies and different cellular responses to the presence of thymidine analogs^24,25^. For the trypanosomatids studied here, species-specific characteristics (such as doubling time) possibly shape the cellular responses to BrdU/EdU incorporation. *T. cruzi* has a doubling time estimated at 24 h^9^, while *T. brucei* and *L. major* have a doubling time estimated at ∼8 h^9,25^. These differences could drastically impact the proportion of BrdU/EdU-positive cells after 24 h of exposure, as well as after 72 h of recovery from BrdU/EdU exposure. For instance, if we consider that the cells are in continuous growth and that BrdU/EdU incorporation occurs consistently without altering the doubling time or proliferative capacity, it would be expected that, after 24 h of exposure, *T. cruzi*, *T. brucei* and *L. major* would show 100% BrdU/EdU-labeled cells (Supplementary Fig. 3). Nevertheless, following a 72 h of recovery period after BrdU/EdU exposure, it would be expected that *T. cruzi* would show 12.5% BrdU/EdU-labeled cells, while *T. brucei* and *L. major* would likewise show 1.37% BrdU/EdU-labeled cells (Supplementary Fig. 3). Although this is merely a hypothetical prediction based on an ideal non-existent environment, the values we measured for the corresponding periods, especially after 72 h of recovery, are far from those predicted (Fig. 3), evidencing that BrdU/EdU incorporation alters the normal behavior of trypanosomatids.

BrdU has already been demonstrated as an agent capable of generating an increase in Sister-Chromatid Exchanges (SCEs) in CHO-K cells^37^, endoreduplication in AA8 cells^38^, accumulation of cells in G1 and imbalance of the dNTP pool in yeast^39^, and an increase in the HPRT mutational rate in CHO cells^22^. Despite these alterations evidenced for several cell types, the effects caused by BrdU in *T. cruzi*, *T. brucei* and *L. major* were relatively mild, maintaining the cells viable and with good proliferation capacity (Fig. 1 and 2). Although we observed a slight increase in the sub-G1 population for *T. brucei* after the 72 h of recovery period (Fig. 2) and in DNA damage signaling (increased γH2A intensity) for all the three trypanosoamatids (Fig. 4), these parasites seem to tolerate the presence of BrdU, despite the potential for the induction of structural changes based on our *in silico* data (Fig. 6; Supplementary Fig. 2). Further studies are needed to investigate whether BrdU indeed causes DNA alterations when incorporated into trypanosomatids genome. If so, we might wonder whether these organisms would be able to repair BrdU-generated DNA lesions quickly and efficiently^40–42^, which would explain the mild effects observed. If BrdU does not cause sufficient alterations to trypanosomatid DNA to trigger a DNA damage response, this could also explain the observed behavior.

In contrast, our findings involving EdU align with the results observed by other research groups in different cell types. As a result of EdU incorporation, researchers have been observing a reduction in cell viability at concentrations higher than 7 µM^20^, cell cycle arrest in the first DNA replicative cycle^43^, chromosomal aberrations^22^, increased DNA damage signaling (increased γH2AX signal)^22^, increased SCE formation^22^, reduction in cell viability^22,44^, delayed cell cycle progression^44^, and apoptosis^44^.

This set of results is in consonance with what we observed in *T. cruzi*. This parasite showed proliferation impairment and reduction in cell viability in response to 50 and 100 μM EdU (Fig. 1A, B). Moreover, the DNA content profile of EdU-treated groups showed a significant accumulation of cells in S (Fig. 2A, B), most likely due to EdU persistence (Fig. 3A, B), and consequent DNA damage signaling evidenced by the presence of a high proportion of γH2A-positive cells (Fig. 4A-C). However, after 72 h of recovery from EdU exposure (day 5), the percentage of *T. cruzi* EdU-incorporated cells remains virtually the same as on day 2 (Fig. 3A, B), suggesting that this parasite lacks efficient mechanisms for removing EdU from its genome, even with intense γH2A signaling (Fig. 4A-C). Another possibility is that *T. cruzi* is in a dormancy-like state (quiescence), a phenomenon previously reported in response to different types of stress^19,45^. However, quiescent cells (G0 state) typically exhibit a DNA content equivalent to the G1 phase, as they are not committed to the cell cycle^46^. We did not observe *T. cruzi* cells accumulating in the G1 phase/G0 state after EdU exposure. Instead, our data suggest that EdU promotes the accumulation of *T. cruzi* cells in the S phase (Fig. 2A, B), possibly due to replication stress. Studies have suggested that *T. cruzi* exhibits similar phenotypes in response to various types of DNA damage^47–49^, which supports that EdU, at concentrations of 50 and 100 µM and with an exposure period of 1 to 24 h, indeed causes substantial DNA damage in this parasite. However, further studies are needed to specifically classify the type of DNA damage predominantly induced by the presence of EdU in *T. cruzi*. Moreover, it is worth highlighting that we conducted our experiments using *T. cruzi* epimastigote (insect-stage) forms, which may restrict the scope of our results, since amastigote (mammal-stage) forms were not analyzed.

*T. brucei*, in contrast, did not behave like *T. cruzi* and appeared to be indifferent to the presence of EdU (up to 100 μM) (Figs. 1C, 1D, 2C, 2D). Interestingly, this thymidine analog only generated a slight increase in γH2A signal intensity (Fig. 4D-F). A possible explanation for this peculiar behavior of *T. brucei* is that this parasite is tolerant to the DNA damage caused by EdU incorporation and apparently possesses molecular mechanisms to continue DNA replication even in the presence of DNA damage, a behavior already proposed, in another context^50^. Apparently, *T. brucei* lacks a strict cell cycle checkpoint. Unlike other trypanosomatids, DNA damage in this organism does not trigger a rigorous checkpoint to halt the cell cycle^50,51^. This high tolerance for DNA damage facilitates the generation of genetic diversity, particularly in subtelomeric regions, which aids in antigenic variation and evasion of the host’s immune system^52^. However, it is worth noting that our results were limited to the evaluation of *T. brucei* procyclic (insect-stage) forms, and it remains unknown whether this behavior would also be observed in the bloodstream (mammal-stage) forms, as already observed for other types of DNA damage agents^53^. We also cannot ignore the fact that the population of *T. brucei* used in our experiments naturally exhibited (in the control groups) a higher proportion of γH2A-positive cells (38,58% ±3,56 – Fig. 4E) than that observed in another study (∼25%)^51^. These data suggest that our cells had a high basal level of persistent DNA damage foci, whose origin remains unknown.

For *L. major*, the effects of EdU were rapid and severe for part of the population, both in terms of decreasing parasite density and viability (Fig. 1E, F), as well as in altering the DNA content profile (Fig. 2E, F). Our data suggest that part of the *L. major* population dies following EdU exposure, and the surviving population resumes a vigorous proliferation and viability 72 h after recovery (Fig. 1E, F). A similar behavior has already been observed in *L. amazonensis*, where H_2_O_2_-induced genotoxic stress appeared to select for the fittest parasites^54^. However, we did not re-expose the surviving *L. major* groups that showed high viability on day 5 (72 h after exposure to 50 and 100 µM EdU) to determine whether they would be more resistant to a subsequent round of EdU incorporation. Furthermore, another study showed that when promastigote forms of *L. mexicana* are exposed to hydroxyurea (a replication stress inducer), a portion of the population dies via apoptosis^55^. However, the cells that survive enter a state of cell cycle arrest^55^, possibly to activate DNA repair pathways and resume proliferation later on^56^. Interestingly, another study demonstrates that the presence of apoptotic parasites in a virulent inoculum of *Leishmania* promastigotes is of paramount importance to enable the intracellular survival of the viable parasites^57^. Similar behavior is also observed when amastigote forms of *Leishmania* are exposed to drugs such as pentavalent antimonials, miltefosine, and amphotericin B^56,58^. Together, these studies corroborate our findings and suggest that the death of part of the *Leishmania* population may act as a form of ‘cellular altruism’^57,59,60^, wherein the release of vesicles and other factors by the dead cells could diminish the intensity of the host’s immune response while simultaneously stimulating vigorous proliferation among the survivors^56–59^. However, as in *Trypanosoma* spp., our study was limited to evaluating *L. major* promastigote (insect-stage) forms, and we have no evidence that this behavior in response to EdU incorporation also occurs in amastigote (mammal-stage) forms.

In general, most altered phenotypes for all trypanosomatids analyzed here were induced by the presence of EdU in their genomes. A recent study suggests that the presence of EdU in DNA can lead to possible stalling or collapse of the replication fork^22^. Furthermore, other studies have proposed that the presence of EdU could generate ICLs in mammalian cells^20,21^. Since ICLs covalently link the two complementary DNA strands^61^, these structures can act as a physical barrier to the CMG complex, being responsible for both the stalling and the collapse of replication forks^62^. ICLs also may generate discrepancies in the number of H-bonds^63^, something that was observed in our molecular dynamic simulations due to the presence of EdU (Fig. 6). Furthermore, the ICLs could trigger a DNA damage response involving the TC-NER pathway to remove EdU from the genome, as recently proposed^23^. Whether EdU indeed causes ICLs in trypanosomatids, and whether these parasites are capable of repairing such lesions via the TC-NER pathway, remain open questions that require further investigation.

Taken together, our results indicate that 10 µM EdU is a safe concentration for the three trypanosomatids analyzed here, although it may not be sufficient for efficient detection. At the other concentrations tested (50 and 100 µM), EdU exhibited higher genotoxic potential than BrdU, and *T. cruzi*, *T. brucei*, and *L. major* respond markedly differently to the presence of this analog, showing significant differences regarding viability, cell cycle progression, analog retention time, and DNA damage signaling. In summary, our findings suggest that using the thymidine analogs to monitor DNA synthesis in trypanosomatids should be done with caution, especially when using EdU. The objective of the assay must be analyzed carefully, and the BrdU/EdU concentrations adjusted according to the required incorporation period and cellular characteristics, such as doubling time. Otherwise, the data obtained may be biased by DNA damage response caused by the incorporation of the respective thymidine analog, not only in trypanosomatids but in any cell type. We believe this study may serve as an atlas to guide the initial selection of thymidine analog concentrations and exposure periods.

## Material and Methods

### Trypanosomatid cultures and growth curves

*T. cruzi* epimastigote forms (CL Brener strain) were cultured at 27 °C in liver-infusion tryptose (LIT) medium supplemented with 10% heat-inactivated fetal bovine serum (FBS) (Cultilab), 10 μg.mL^-1^ Hemine (Sigma-aldrich), and 1% (v/v) antibiotic solution (streptomycin/penicillin). *T. brucei* trypomastigotes (procyclic forms) (TREU927 strain) were cultured at 27 °C in SDM79 medium supplemented with 10% (v/v) heat-inactivated FBS and 1% (v/v) antibiotic solution (streptomycin/penicillin). *L. major* promastigotes (Friedlin strain) were cultured at 27 °C in M199 medium (Thermo-Fischer Scientific) supplemented with 10% (v/v) heat-inactivated FBS, 25 mM HEPES (USB corporation), 0.1 mM Adenine (Sigma-Aldrich), 0.0001% Biotin (Sigma-Aldrich), and 1% (v/v) antibiotic/antimycotic solution.

For the growth curves, *T. cruzi* epimastigotes and *T. brucei* trypomastigotes were started at a concentration of 2.10^6^ cells.mL^-1^ and counted daily for 5 days. *L. major* promastigotes were started at a concentration of 1.10^6^ cells.mL^-1^ and counted daily for 5 days. The counting was done using a Neubauer chamber (Optik Labor). The final concentration was calculated using the following formula: *x̄ quadrants dilution^−1^*·10^4^. The standard deviation (SD) in each growth curve was calculated from three independent experiments.

### Viability assay using 3-(4,5-dimethylthiazol-2-yl)-2,5-diphenyltetrazolium bromide (MTT)

Viability assays were carried out using 3-(4,5-dimethylthiazol-2-yl)-2,5-diphenyltetrazolium bromide (MTT), a compound that is reduced by NAD(P)H-dependent cellular oxidoreductases to a purple formazan compound in living cells^64^. Briefly, the same volume (1–2 mL, depending on cell density) from each trypanosomatid species culture was harvested at specific points (days) in the growth curve and placed into a 48-well flat-bottom plate. Next, the samples containing parasites were centrifuged (1000 × *g* for 5 min) and the medium replaced with 3 mg.mL^-1^ MTT solution diluted in PBS (137 mM NaCl, 2.7 mM KCl, 10 mM Na_2_HPO_4_ and 2 mM KH_2_PO_4_, pH 7.4). The parasites were then resuspended and incubated for 4 h with gentle shaking. Next, the samples were centrifuged (1000 × *g* for 5 min), MTT solution discarded, and the resultant formazan crystals solubilized with 100% dimethyl sulfoxide (DMSO). The absorbance for each well was measured at 570 nm using a microplate spectrophotometer (Epoch – Biotek). The absorbance values obtained were used to infer the viability percentage relative to the control group (non-treated) of each trypanosomatid species analyzed. The SEM (standard error of the mean) in each bar graph was calculated from three independent experiments, each of which was performed as a technical triplicate.

### Flow cytometry analysis

For DNA content analysis, each trypanosomatid species was harvested, centrifuged (1000 × *g* for 5 min), washed twice using PBS, and fixed/permeabilized with 70% cold ethanol overnight at 4 °C. The pellet containing parasites was washed once with 10 mL of PBS and resuspended in 500 μL of a solution containing 10 μg.mL^-1^ of Propidium Iodide (PI) and 100 μg.mL^-1^ of RNase A (diluted in PBS). After 45 min of incubation at 37 °C, the parasites were analyzed by flow cytometry using a CytoFLEX LX (Beckman Coulter). The FCS files generated were analyzed using both FlowJo vX.0.7 (Tree Star) and Floreada.io (https://floreada.io/). The SEM (standard error of the mean) in each bar graph of the cell cycle analyses was calculated from three independent experiments.

### Immunofluorescence assays (IFA)

For the detection of BrdU-incorporated cells in the different periods analyzed, we used α-BrdU antibodies (rabbit – Thermo Fischer; or mouse – DB Biosciences) and followed a protocol previously described^9^, with the following modifications. Briefly, each group of each trypanosomatid analyzed was harvested, centrifuged (1000 × *g* for 5 min), washed twice using PBS, fixed for 10 min with 4% sterile paraformaldehyde (Merck) diluted in PBS, and added to a 0.1% poly-L-lysine pre-treated slide (Knittel Glaser). The slides were then permeabilized with 0.1% Triton X-100 diluted in PBS for 5 min and briefly washed with PBS. Next, the DNA was denatured in a 3 M HCl solution for 20 min, then neutralized using 40 μL of 0.1 M borate buffer (100 mM H_3_BO_3_, 75 mM NaCl, 25 mM Na_2_B_4_O_7_, pH 7.5) for 10 min. After this step, the slides were washed three times with PBS then allowed to dry. Afterward, α-BrdU (rabbit – Thermo Fischer; or mouse – DB Biosciences) was diluted 1:500 in 4% BSA solution, added to the slide and the slide was incubated overnight at 4 °C. Next, the slides were carefully washed with PBS and the secondary antibody (Alexa Fluor 488 anti-rabbit IgG – Thermo Fischer or Alexa Fluor 594 anti-mouse IgG – Thermo Fischer, diluted 1:1000 in 4% BSA) was added and incubated for 2 h at 4 °C (protected from light). Then, the slides were washed carefully with PBS and left to dry momentarily. Vectashield containing DAPI (VectorLabs) or DAPI fluoromount G (ThermoFischer) was used as an anti-fade mounting solution, and to stain nuclei and kinetoplast DNA. Images were acquired on a Zeiss Axiovert 200 M fluorescent microscope fitted with an HXP 120 V lamp (Zeiss) using a 100x oil lens or on a Leica Stellaris 8 fluorescent microscope fitted with a 100x oil lens. When necessary, the images were superimposed and processed using ImageJ (1.46r) software^65^.

For the DNA damage analyses in trypanosomatids, an α-γH2A antibody whose affinity for trypanosomatid species was validated in other studies^19,26,27^ was used. The antibody was gifted by Professor Richard McCulloch from University of Glasgow, UK. Briefly, parasites were harvested, centrifuged (1000 × *g* for 5 min) to remove the medium and processed as described above. α-γH2A antiserum (rabbit) was diluted 1:500 in 4% BSA solution, added to the slide and incubated overnight at 4 °C. After, the slides were carefully washed with PBS, then incubated with the secondary antibody (Alexa Fluor 647 anti-rabbit IgG – Thermo Fischer, diluted in 4% BSA – 1:1000) for 2 h at 4 °C (protected from light). Slides were then washed with PBS and left to dry momentarily. Slides were mounted using Vectashield containing DAPI (VectorLabs) to stain nuclei and kinetoplast DNA. Images were captured on a Zeiss Axiovert 200 M fluorescent microscope fitted with an HXP 120 V lamp (Zeiss) using a 100x oil lens. When necessary, the images were superimposed and processed using ImageJ (1.46r) software^65^.

### ‘Click’ chemistry reaction for EdU detection

For the detection of EdU-incorporated cells at 1 h exposure, 24 h exposure, and 72 h post-EdU recovery, we used ‘click’ chemistry reactions as previously described^9^, with the following modifications. Briefly, each trypanosomatid species was harvested, centrifuged (1000 × *g* for 5 min), washed twice using PBS, fixed for 10 min with 4% sterile paraformaldehyde (Merck) diluted in PBS, and added into a 0.1% poly-L-lysine pre-treated slide (Knittel Glaser). The parasites were then permeabilized using 0.1% Triton X-100 diluted in PBS for 5 min and briefly washed with PBS. To detect incorporated EdU, we used 50 µL of Click-EdU detection solution (3 mM CuSO_4_, 100 mM C_6_H_8_O_6_, and 5 mM Alexa fluor azide 488 – Thermo Fischer, diluted in PBS) for 1 h (protected from light). Finally, the cells were washed twice with PBS and the slides mounted using Vectashield containing DAPI (VectorLabs) to stain nuclei and kinetoplast DNA. Images were acquired as described above. When necessary, the images were superimposed and processed using ImageJ (1.46r) software^65^.

### Chemical characteristics of dT, BrdU and EdU

*In silico* predictions of the chemical characteristics of the nucleosides dT (deoxythymidine), BrdU (5-Bromo-2’-deoxyuridine), and EdU (5-Ethynyl-2’-deoxyuridine) were performed as follows. Briefly, ACD/ChemSketch 2023.2.4 was used to generate the 3D structures of dT, BrdU and EdU with the addition of hydrogens, followed by optimization as a preliminary step. Next, using Avogadro 1.2.0, we performed a new optimization using the Merck Molecular Force Field (MMFF94), a program ideal for small organic molecules^66^. For the conformational analysis, we chose the Systematic Rotor Search technique, which explores all possible combinations for the torsion angles of each rotatable bond to find the lowest energy conformation^67^. Using GaussView 5.0.8, we selected the optimization and frequency calculations using the Density Functional Theory (DFT) method – B3LYP/6-31G. Using Gaussian09 package^68^, we generated the outputs and set calculations to obtain the structures in their minimum energy state. Using Gaussian09 package, we estimated the Highest Occupied Molecular Orbital (HOMO), Lowest Unoccupied Molecular Orbital (LUMO), and the partial charge of the atoms. Finally, using GaussView 5.0.8, we generated the molecular electrostatic potential (MEP) maps for each nucleoside analyzed using the total density parameters. The values for HOMO-LUMO gap, electronegativity, and electrophilicity were estimated using the Gaussian09 package^68^.

### Molecular dynamics of DNA molecules after BrdU/EdU incorporation

To perform molecular dynamics simulations of DNA molecules, DNA was first represented using a specialized all-atom molecular dynamics (MD) parameter set called Amber OL15 force field^69^. EdU and BrdU were optimized using the Gaussian09 package using a B3LYP algorithm^68^. The optimized structures were used for calculating partial charges utilizing a Hartree-Fock algorithm. The residual (resp) charges were calculated with the antechamber24 package^70^. Partial atomic charges, bonded, and non-bonded parameters were added to the force field files. The topology was generated using the *pdb2gmx* module from GROMACS 2022.4 package^71,72^.

For simulations, five dsDNA molecules (20 mer) were analyzed: a control DNA molecule (BrdU/EdU non-incorporated), a DNA with BrdU incorporated in one strand, a DNA with BrdU incorporated in both strands, a DNA with EdU incorporated in one strand, and a DNA with EdU incorporated in both strands (Fig. 6A). EdU and BrdU bases were distributed next to each possible DNA base (A, C, or G), in the same and/or in the complementary strand, to cover the maximum number of possible combinations. The double helixes structures were built in the initial standard conformations using the Avogadro package^73^. Each of the five structures was centered in an octahedral box, with a side length of 8 nm and solvated with 12411 to 12415 TIP3P water molecules^74^, 74 Na^+^ and 36 Cl^−^ ions (0.15 M NaCl) to maintain electroneutrality^75^. Energy minimization was performed using a Steepest Gradient Descent algorithm with a maximum force of 1 kJ. We used three 1 ns sequential NVT equilibration steps, warming the system from 200 K to 300.15 K (30 °C) with 10 KJ position restraints on the DNA molecule. At 300.15 K, eight 1 ns NVT steps were performed, reducing the position of the restraint force at each step. The NPT equilibration was performed for 1 ns without position restraints. The resulting systems were simulated for 500 ns in three parallel calculations for each system.

Simulations were carried out using GROMACS 2022.4 package^71^. The simulations used a 2 fs time step with a Leap-Frog integrator^76^. H-bonds were restrained with the P-LINCS algorithm^77^. We linearly removed the center of mass for the entire system every 100 simulation steps. The electrostatic interactions were treated using the Particle Mesh Ewald (PME) method^78^, with a real-space cutoff radius of 0.9 nm. Van der Waals interactions were treated with a 0.9 nm cutoff radius using a 0.9 nm r-list radius. The temperature was maintained at 300.15 K using a Velocity Rescale thermostat^79^, with a coupling constant of 1.0 ps, and DNA, water, and ions were coupled in two groups. The pressure was kept isotropically at 1.013 bar using a Parrinello-Rahman barostat^80^ with a 10.0 ps coupling constant and a compressibility of 4.5×10^-5^.

RMSD and RMSF analyses were conducted for the DNA backbone, using the initial structure as a reference. Backbone dihedral angles were computed individually for each dihedral type. H-bonds between chains were computed for each base. The structures visited from 100 ns to 500 ns of simulation were averaged in clusters. Each cluster represented the average of structures with an RMSD between them lower than a cutoff value. The clusters were ranked by probability. For control DNA, a 3 Å cutoff was used for clustering, while a 2 Å cutoff was used for the DNAs containing EdU and BrdU. The number of H-bonds, and the RMSD and RMSF values were averaged after 100 ns of simulation. For each simulation system, the moving averages of the three replicas were calculated, and the pooled standard deviation was used to determine the moving standard deviations and error bars. Python was used for data analysis and plotting, utilizing the NumPy^81^, Pandas, and Matplotlib packages. Independent *t*-tests were performed with the SciPy package^82^.

### Statistics and reproducibility

Statistical analyses were performed using GraphPad Prism 9.0.0 package (GraphPad Software, Inc.) unless otherwise stated. Normal distribution was assessed for all datasets using a Shapiro–Wilk normality test. Significant differences between groups were identified using two-tailed Student’s t-test (paired or unpaired, as appropriate). Comparisons of more than two groups were performed with one-way ANOVA followed by Student–Newman–Keuls test. Data are represented as mean ± SEM (standard error of the mean), unless otherwise stated.

## Data availability

The original figures, graphs and dataset that support the findings of this study are available from the corresponding author upon reasonable request.

## Acknowledgments

The authors are grateful to Marcelo Nunes Silva for his immeasurable help in the preparation of reagents/solutions and laboratory organization. We would also like to thank Professors Maurício Baptista (University of São Paulo, Brazil), Alexander Henning Ulrich (University of São Paulo, Brazil), Sérgio Schenkman (Federal University of São Paulo), and Luiz R. O. Tosi (University of São Paulo, Brazil), for making their reagents, equipment, and laboratory available for our use.

