## Supplementary Material for "A comparative analysis of BrdU and EdU genotoxicity shows that trypanosomatids exhibit unique responses to the presence of these thymidine analogs in their DNA"

[1]Department of Biochemistry, Institute of Chemistry, University of São Paulo (USP), São Paulo, SP, Brazil; [2]Department of Genetics, Microbiology and Immunology, Biosciences Institute, São Paulo State University (UNESP), Botucatu, SP, Brazil; [3]Department of Cellular and Molecular Biology, Ribeirão Preto Medical School, University of São Paulo, Ribeirão Preto, 14040-900 SP, Brazil; [4]University of Glasgow Centre for Parasitology, School of Infection and Immunity, Sir Graeme Davies Building, 120 University Place, Glasgow G12 8TA, United Kingdom.

[#] **These authors contributed equally to this work**

***Corresponding author**: Marcelo S. da Silva – 748, Professor Lineu Prestes Avenue, block 9, room 916, São Paulo, SP, Brazil. Postal code: 05508-000. phone +55 (11) 3091 2696;

**Supplementary Figure 1**

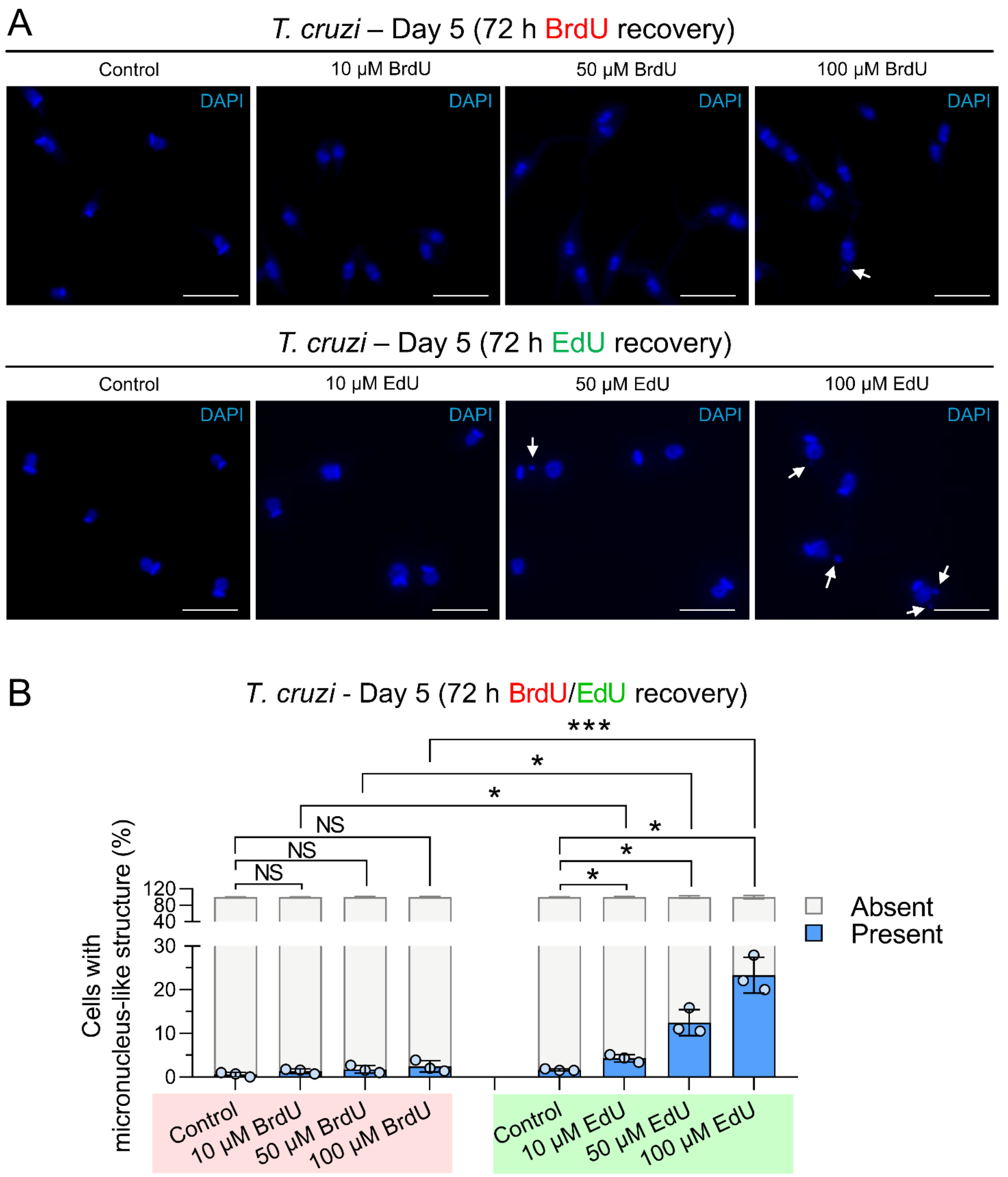


**Supplementary Fig. 1. EdU exposure generates DNA-containing micronucleus-like structures in *T. cruzi*. A**. Representative images of the DAPI-stained *T. cruzi* cells after 72 h of recovery from BrdU/EdU exposure. White arrows point to DNA-containing micronucleus-like structures. The bar represents 10 μm. **B**. Quantification of the proportion of *T. cruzi* cells presenting DNA-containing micronucleus-like structures. The values and the standard error of the mean (SEM) were obtained from assays performed in triplicate. Statistical significance was assessed using unpaired two-sided Student's t tests. Significance is indicated as: *$p < 0.05$, **$p < 0.01$, ***$p < 0.001$. NS denotes no significant difference according to Student's *t*-test.

**Supplementary Figure 2**

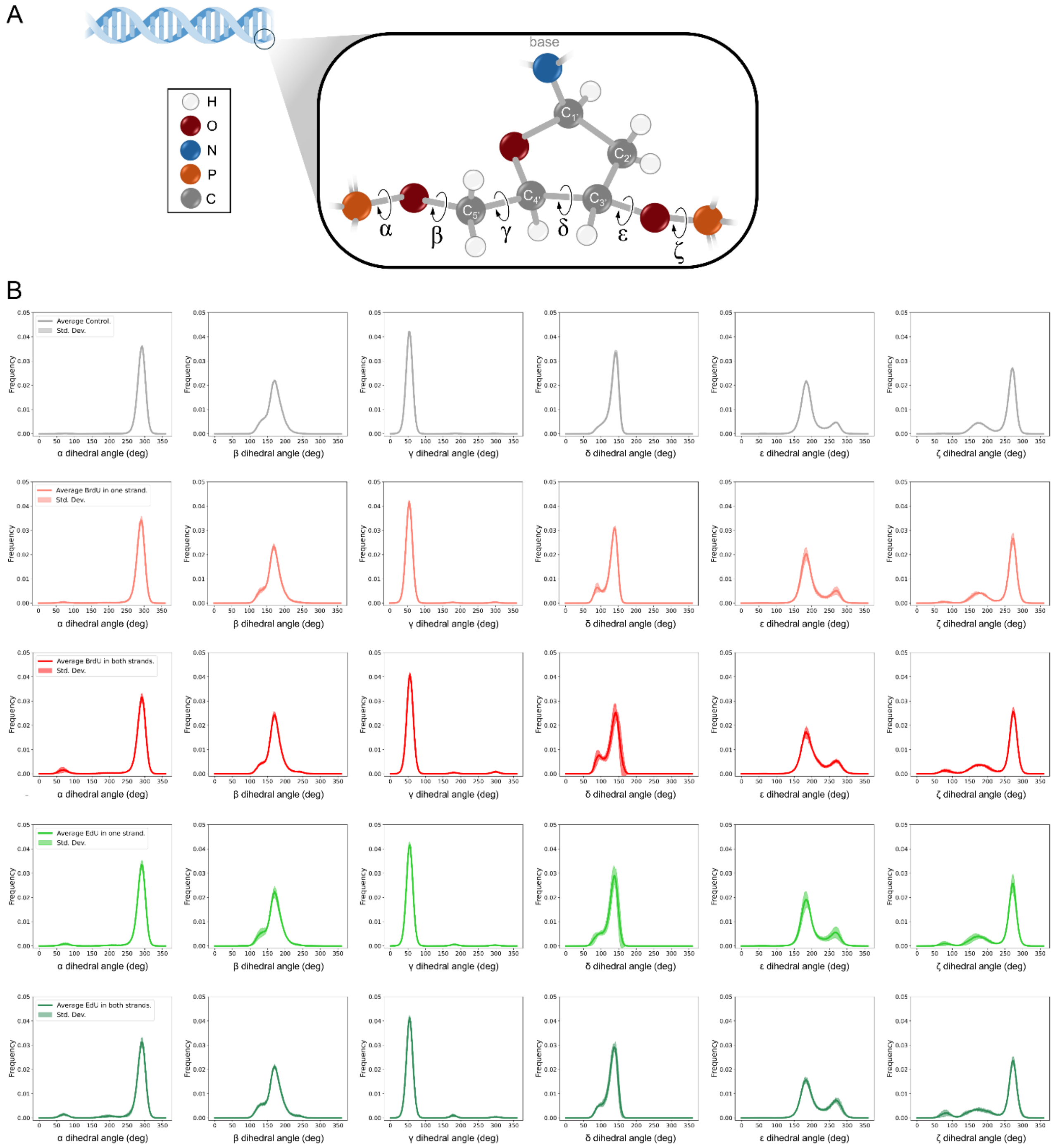


**Supplementary Fig 2. Frequency of dihedral angles of DNA structure generated during molecular dynamics simulations**. **A**. Schematic representation of the positions of dihedral angles in the DNA structure. **B**. Frequency of dihedral angles obtained during molecular dynamics simulations. The Y-axis represents the dihedral angle frequency, and the X-axis represents the values in degrees (deg). Gray histograms represent the control DNA (BrdU/EdU non-incorporated DNA); light red indicates the simulation with BrdU in one strand, dark red in two strands; light green indicates the simulation with EdU on one strand and dark green on two strands of DNA.

### Supplementary Figure 3

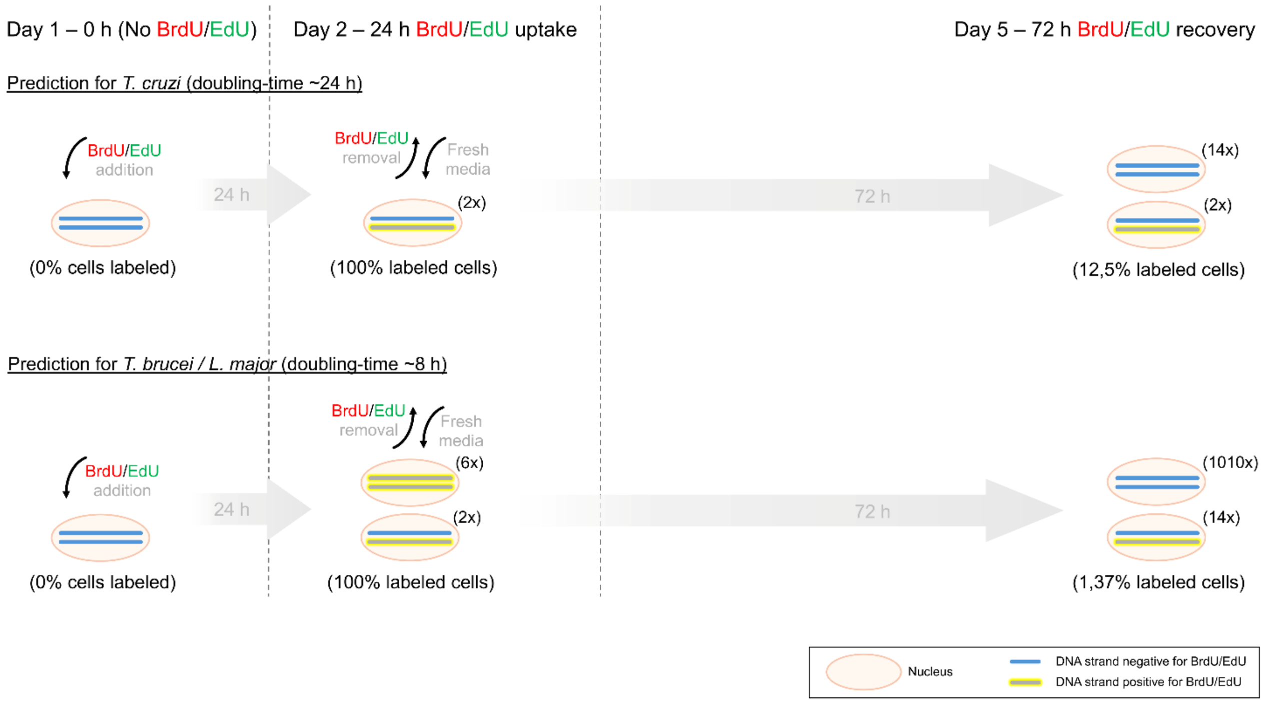


**Supplementary Fig 3. Theoretical prediction of the proportion of labeled cells after 24 h BrdU/EdU exposure and after 72 h of recovery following exposure**. For this estimative, we considered that all trypanosomatids analyzed grew continuously (without any replication impairment). We also used a doubling time of 24 h for *T. cruzi* and considered 8 h as a doubling time for *T. brucei* and *L. major*. The relative number of cells/nuclei generated with that phenotype after the indicated period is shown in parentheses.

**Supplementary Table 1.** Numerical sources (mean and standard deviation) of the *Trypanosoma cruzi* cell cycle analysis that were presented in Figure 2A, B.

**A.** Day 1 (1 h BrdU uptake)

| | Sub G1 (%) | G0/G1 (%) | S (%) | G2/M (%) | >G2/M (%) |
|---|---|---|---|---|---|
| Control | 1.95 ±0.54 | 31.93 ±1.56 | 19.53 ±1.09 | 35,33 ±2.09 | 10.18 ±1.88 |
| 10 µM BrdU | 2.68 ±0.34 | 32.67 ±0.18 | 22.33 ±0.71 | 31,67 ±0.69 | 9.19 ±0.49 |
| 50 µM BrdU | 2.92 ±0.32 | 33.03 ±0.62 | 21.23 ±0.42 | 33,50 ±1.07 | 8.04 ±0.58 |
| 100 µM BrdU | 2.77 ±0.8 | 34.5 ±0.53 | 23.03 ±1.56 | 32,20 ±2.13 | 6.59 ±0.65 |

**B.** Day 2 (24 h BrdU uptake)

| | Sub G1 (%) | G0/G1 (%) | S (%) | G2/M (%) | >G2/M (%) |
|---|---|---|---|---|---|
| Control | 2.06 ±0.11 | 32.33 ±0.51 | 22.50 ±0.2 | 34.07 ±0.29 | 8.88 ±0.64 |
| 10 µM BrdU | 1.95 ±0.05 | 31.97 ±0.64 | 21.83 ±0.89 | 34.27 ±0.31 | 9.98 ±0.29 |
| 50 µM BrdU | 2.18 ±0.2 | 31.13 ±1.11 | 22.97 ±0.56 | 34.47 ±1.09 | 9.42 ±0.63 |
| 100 µM BrdU | 2.23 ±0.29 | 31.47 ±1.02 | 23.00 ±0.87 | 34.50 ±0.27 | 8.74 ±0.49 |

**C.** Day 5 (72 h BrdU recovery)

| | Sub G1 (%) | G0/G1 (%) | S (%) | G2/M (%) | >G2/M (%) |
|---|---|---|---|---|---|
| Control | 1.28 ±0.15 | 43.77 ±0.24 | 13.77 ±0.18 | 29.17 ±0.51 | 12.20 ±1.07 |
| 10 µM BrdU | 2.06 ±0.54 | 41.90 ±1.13 | 15.23 ±0.49 | 27.80 ±0.33 | 13.00 ±0.53 |
| 50 µM BrdU | 1.77 ±0.14 | 43.03 ±0.22 | 15.43 ±0.69 | 28.50 ±0.6 | 11.33 ±0.58 |
| 100 µM BrdU | 1.49 ±0.29 | 42.83 ±0.49 | 16.17 ±0.69 | 27.07 ±0.51 | 12.60 ±0.93 |

**D.** Day 1 (1 h EdU uptake)

| | Sub G1 (%) | G0/G1 (%) | S (%) | G2/M (%) | >G2/M (%) |
|---|---|---|---|---|---|
| Control | 1.49 ±0.05 | 33.93 ±0.44 | 16.73 ±0.96 | 38.83 ±1.62 | 9.17 ±0.32 |
| 10 µM EdU | 1.57 ±0.16 | 34.60 ±0.33 | 16.33 ±0.69 | 36.87 ±0.49 | 10.80 ±0.8 |
| 50 µM EdU | 1.57 ±0.18 | 35.43 ±0.51 | 15.83 ±0.51 | 37.37 ±0.91 | 9.97 ±0.22 |
| 100 µM EdU | 1.63 ±0.21 | 36.53 ±0.38 | 16.77 ±0.42 | 35.83 ±1.09 | 9.33 ±0.63 |

**E.** Day 2 (24 h EdU uptake)

| | Sub G1 (%) | G0/G1 (%) | S (%) | G2/M (%) | >G2/M (%) |
|---|---|---|---|---|---|
| Control | 1.60 ±0.11 | 33.67 ±0.49 | 20.93 ±0.29 | 37.27 ±0.56 | 6.86 ±0.28 |
| 10 µM EdU | 1.81 ±0.18 | 32.97 ±0.44 | 23.23 ±0.78 | 35.83 ±0.89 | 6.19 ±0.45 |
| 50 µM EdU | 1.68 ±0.09 | 33.27 ±0.49 | 21.13 ±0.56 | 38.17 ±0.11 | 5.87 ±0.25 |
| 100 µM EdU | 1.74 ±0.09 | 33.03 ±0.78 | 21.03 ±0.38 | 39.53 ±0.58 | 4.85 ±0.25 |

**F.** Day 5 (72 h EdU recovery)

| | Sub G1 (%) | G0/G1 (%) | S (%) | G2/M (%) | >G2/M (%) |
|---|---|---|---|---|---|
| Control | 2.47 ±0.48 | 42.67 ±0.18 | 15.50 ±0.4 | 27.53 ±0.31 | 11.80 ±0.33 |
| 10 µM EdU | 5.33 ±0.09 | 24.25 ±0.43 | 18.95 ±0.63 | 38.70 ±0.07 | 12.85 ±0.1 |
| 50 µM EdU | 4.17 ±0.13 | 14.23 ±0.51 | 34.57 ±0.82 | 39.77 ±1.22 | 6.88 ±0.35 |
| 100 µM EdU | 4.15 ±0.6 | 10.70 ±0.33 | 38.10 ±1.2 | 40.27 ±1.76 | 6.76 ±0.47 |

**Supplementary Table 2** – Numerical sources (mean and standard deviation) of the *Trypanosoma brucei* cell cycle analysis that were presented in Figure 2C, D.

**A.** Day 1 (1 h BrdU uptake)

| | Sub G1 (%) | G0/G1 (%) | S (%) | G2/M (%) | >G2/M (%) |
|---|---|---|---|---|---|
| Control | 4.48 ±0.59 | 61.45 ±0.73 | 11.27 ±0.82 | 19.2 ±0.87 | 2.36 ±0.52 |
| 10 μM BrdU | 4.4 ±0.57 | 62.13 ±0.22 | 11.18 ±0.16 | 18.77 ±0.71 | 2.61 ±0.16 |
| 50 μM BrdU | 4.56 ±0.59 | 62.27 ±0.91 | 11.57 ±0.31 | 18.73 ±0.22 | 2.02 ±0.16 |
| 100 μM BrdU | 4.58 ±0.69 | 62.97 ±0.49 | 11.1 ±0.33 | 19.2 ±0.33 | 2.1 ±0.18 |

**B.** Day 2 (24 h BrdU uptake)

| | Sub G1 (%) | G0/G1 (%) | S (%) | G2/M (%) | >G2/M (%) |
|---|---|---|---|---|---|
| Control | 2.92 ±0.1 | 71.97 ±0.51 | 7.18 ±0.21 | 14.27 ±0.58 | 2.42 ±0.25 |
| 10 μM BrdU | 3.65 ±0.29 | 72.63 ±0.58 | 7.63 ±0.42 | 13.6 ±0.53 | 1.99 ±0.21 |
| 50 μM BrdU | 3.6 ±0.2 | 72.5 ±0.53 | 6.95 ±0.06 | 14.13 ±0.31 | 1.72 ±0.12 |
| 100 μM BrdU | 3.57 ±0.33 | 72.43 ±0.51 | 7.27 ±0.57 | 14.63 ±0.24 | 2.0 ±0.09 |

**C.** Day 5 (72 h BrdU recovery)

| | Sub G1 (%) | G0/G1 (%) | S (%) | G2/M (%) | >G2/M (%) |
|---|---|---|---|---|---|
| Control | 4.91 ±0.25 | 78.07 ±1.02 | 1.7 ±0.32 | 12.20 ±0.2 | 2.92 ±0.13 |
| 10 μM BrdU | 8.48 ±1.83 | 74.33 ±3.02 | 1.7 ±0.29 | 12.7 ±0.73 | 2.4 ±0.3 |
| 50 μM BrdU | 10.14 ±1.78 | 72.6 ±3.13 | 2.07 ±0.4 | 12.73 ±0.78 | 3.22 ±0.28 |
| 100 μM BrdU | 12.97 ±0.62 | 68.93 ±0.71 | 2.02 ±0.23 | 13.59 ±0.34 | 2.57 ±0.21 |

**D.** Day 1 (1 h EdU uptake)

| | Sub G1 (%) | G0/G1 (%) | S (%) | G2/M (%) | >G2/M (%) |
|---|---|---|---|---|---|
| Control | 4.37 ±0.51 | 62.28 ±0.38 | 11.53 ±0.82 | 19.17 ±0.89 | 2.38 ±0.54 |
| 10 μM EdU | 4.61 ±0.39 | 62.03 ±0.82 | 11.98 ±0.54 | 20.37 ±0.76 | 1.91 ±0.3 |
| 50 μM EdU | 4.01 ±0.54 | 61.60 ±0.47 | 10.18 ±1.24 | 21.73 ±1.62 | 2.63 ±0.25 |
| 100 μM EdU | 4.00 ±0.09 | 63.53 ±0.64 | 11.08 ±0.25 | 20.53 ±0.98 | 1.8 ±0.2 |

**E.** Day 2 (24 h EdU uptake)

| | Sub G1 (%) | G0/G1 (%) | S (%) | G2/M (%) | >G2/M (%) |
|---|---|---|---|---|---|
| Control | 2.87 ±0.11 | 72.07 ±0.38 | 6.94 ±0.35 | 14.4 ±0.4 | 2.42 ±0.25 |
| 10 μM EdU | 3.96 ±0.16 | 71.7 ±0.33 | 7.64 ±0.31 | 14.23 ±0.78 | 2.25 ±0.1 |
| 50 μM EdU | 3.97 ±0.2 | 61.8 ±0.33 | 13.23 ±0.29 | 19.23 ±0.62 | 2.22 ±0.2 |
| 100 μM EdU | 4.65 ±0.37 | 50.63 ±0.56 | 19.3 ±1.47 | 23.3 ±1.93 | 2.46 ±0.25 |

**F.** Day 5 (72 h EdU recovery)

| | Sub G1 (%) | G0/G1 (%) | S (%) | G2/M (%) | >G2/M (%) |
|---|---|---|---|---|---|
| Control | 4.91 ±0.25 | 78.92 ±0.74 | 1.7 ±0.32 | 11.68 ±0.49 | 2.88 ±0.15 |
| 10 μM EdU | 3.43 ±0.57 | 77.1 ±0.87 | 2.65 ±0.29 | 13.5 ±0.33 | 3.15 ±0.25 |
| 50 μM EdU | 3.42 ±0.34 | 77.63 ±0.42 | 2.59 ±0.21 | 13.27± 0.31 | 2.71 ±0.06 |
| 100 μM EdU | 3.78 ±0.54 | 78.07 ±1.84 | 2.90 ±0.34 | 12.37 ± 0.71 | 2.70 ±0.37 |

**Supplementary Table 3** – Numerical sources (mean and standard deviation) of the *Leishmania major* cell cycle analysis that were presented in Figure 2E, F.

**A.** Day 1 (1 h BrdU uptake)

| | Sub G1 (%) | G0/G1 (%) | S (%) | G2/M (%) | >G2/M (%) |
|---|---|---|---|---|---|
| Control | 2.21 ±0.09 | 49.43 ±0.29 | 15.67 ±0.38 | 23.83 ±0.64 | 8.92 ±0.39 |
| 10 μM BrdU | 2.57 ±0.14 | 49.9 ±0.6 | 17.87 ±0.62 | 21.1 ±0.47 | 8.64 ±0.31 |
| 50 μM BrdU | 2.62 ±0.17 | 46.7 ±0.6 | 17.9 ±1.53 | 22.27 ±0.84 | 10.6 ±0.2 |
| 100 μM BrdU | 2.43 ±0.05 | 47.47 ±0.29 | 17.67 ±0.82 | 23.6 ±1.2 | 9.03 ±0.29 |

**B.** Day 2 (24 h BrdU uptake)

| | Sub G1 (%) | G0/G1 (%) | S (%) | G2/M (%) | >G2/M (%) |
|---|---|---|---|---|---|
| Control | 2.51 ±0.17 | 54.0 ±1 | 15.83 ±1.11 | 20.4 ±0.73 | 7.24 ±0.48 |
| 10 μM BrdU | 2.6 ±0.16 | 53.27 ±0.38 | 15.23 ±0.76 | 21.57 ±0.64 | 7.34 ±0.21 |
| 50 μM BrdU | 3.62 ±0.45 | 53.03 ±0.56 | 15.73 ±0.11 | 20.53 ±0.69 | 7.2 ±0.72 |
| 100 μM BrdU | 2.72 ±0.16 | 53.77 ±0.84 | 16.8 ±0.67 | 19.87 ±1.11 | 6.93 ±0.66 |

**C.** Day 5 (72 h BrdU recovery)

| | Sub G1 (%) | G0/G1 (%) | S (%) | G2/M (%) | >G2/M (%) |
|---|---|---|---|---|---|
| Control | 8.3 ±0.66 | 62.9 ±2.73 | 5.13 ±0.3 | 16.90 ±0.47 | 6.87 ±1.84 |
| 10 μM BrdU | 8.19 ±0.37 | 62.17 ±0.51 | 6.31 ±0.35 | 17.4 ±0.27 | 6.10 ±0.3 |
| 50 μM BrdU | 8.72 ±0.22 | 64.03 ±1.42 | 7.41 ±0.53 | 15.33 ±0.62 | 4.52 ±0.22 |
| 100 μM BrdU | 8.75 ±0.12 | 63.87 ±0.76 | 6.81 ±0.11 | 15.87 ±0.38 | 4.86 ±0.25 |

**D.** Day 1 (1 h EdU uptake)

| | Sub G1 (%) | G0/G1 (%) | S (%) | G2/M (%) | >G2/M (%) |
|---|---|---|---|---|---|
| Control | 2.33 ±0.06 | 55.27 ±3.62 | 16.3 ±1.33 | 20.2 ±1.53 | 5.9 ±1.28 |
| 10 μM EdU | 2.21 ±0.19 | 56.8 ±0.67 | 16.23 ±0.91 | 18.67 ±0.24 | 6.18 ±0.52 |
| 50 μM EdU | 2.8 ±0.1 | 56.73 ±0.11 | 15.63 ±0.82 | 18.73 ±0.42 | 6.25 ±0.41 |
| 100 μM EdU | 2.41 ±0.1 | 57.83 ±0.29 | 15.8 ±0.0 | 18.37 ±0.04 | 5.81 ±0.2 |

**E.** Day 2 (24 h EdU uptake)

| | Sub G1 (%) | G0/G1 (%) | S (%) | G2/M (%) | >G2/M (%) |
|---|---|---|---|---|---|
| Control | 5.14 ±0.67 | 53.9 ±0.87 | 13.2 ±0.27 | 18.17 ±1.04 | 9.61 ±0.72 |
| 10 μM EdU | 4.45 ±0.84 | 45.57 ±1.78 | 18.5 ±0.27 | 22.1 ±0.4 | 9.55 ±1.97 |
| 50 μM EdU | 6.82 ±0.22 | 37.73 ±0.38 | 21.27 ±1.24 | 26.33 ±1.11 | 8.06 ±0.68 |
| 100 μM EdU | 6.45 ±0.47 | 34.80 ±1.47 | 20.0 ±0.93 | 29.7 ±1.6 | 9.15 ±0.45 |

**F.** Day 5 (72 h EdU recovery)

| | Sub G1 (%) | G0/G1 (%) | S (%) | G2/M (%) | >G2/M (%) |
|---|---|---|---|---|---|
| Control | 5.27 ±0.16 | 73.33 ±0.22 | 5.13 ±0.24 | 13.13 ±0.49 | 3.24 ±0.13 |
| 10 μM EdU | 9.66 ±0.3 | 66.83 ±0.89 | 7.36 ±0.5 | 13.47 ±0.16 | 2.67 ±0.31 |
| 50 μM EdU | 13.5 ±0.93 | 57.87 ±1.02 | 9.24 ±0.61 | 15.2 ±0.87 | 4.21 ±0.34 |
| 100 μM EdU | 14.13 ±0.44 | 56.13 ±0.42 | 9.36 ±0.5 | 15.4 ±0.13 | 4.92 ±0.19 |

**Supplementary Table 4.** Numerical sources (mean and standard deviation) of the BrdU/EdU retention time in *Trypanosoma cruzi* that were presented in Figure 3A, B.

| **A.** Day 1 (1 h BrdU uptake) | | |
|---|---|---|
| | Cells BrdU positive (%) | Cells BrdU negative (%) |
| Control | 0.0 ±0.0 | 100.0 ±0.0 |
| 10 μM BrdU | 1.0 ±0.33 | 99.0 ±0.33 |
| 50 μM BrdU | 5.59 ±1.8 | 94.4 ±1.8 |
| 100 μM BrdU | 31.5 ±7.67 | 68.5 ±7,67 |

| **B.** Day 2 (24 h BrdU uptake) | | |
|---|---|---|
| | Cells BrdU positive (%) | Cells BrdU negative (%) |
| Control | 0.0 ±0.0 | 100.0 ±0.0 |
| 10 μM BrdU | 13.33 ±0.89 | 86.67 ±0.89 |
| 50 μM BrdU | 24.83 ±2.11 | 75.17 ±2.11 |
| 100 μM BrdU | 66.17 ±5.89 | 33.83 ±5.89 |

| **C.** Day 5 (72 h BrdU recovery) | | |
|---|---|---|
| | Cells BrdU positive (%) | Cells BrdU negative (%) |
| Control | 0.0 ±0.0 | 100.0 ±0.0 |
| 10 μM BrdU | 9.33 ±2.78 | 90.67 ±2.78 |
| 50 μM BrdU | 26.17 ±5.22 | 73.83 ±5.22 |
| 100 μM BrdU | 35.73 ±13.51 | 64.27 ±13.51 |

| **D.** Day 1 (1 h EdU uptake) | | |
|---|---|---|
| | Cells EdU positive (%) | Cells EdU negative (%) |
| Control | 0.0 ±0.0 | 100.0 ±0.0 |
| 10 μM EdU | 22.0 ±3.0 | 78.0 ±3.0 |
| 50 μM EdU | 41.17 ±4.56 | 58.83 ±4.56 |
| 100 μM EdU | 43.17 ±6.11 | 56.83 ±6.11 |

| **E.** Day 2 (24 h EdU uptake) | | |
|---|---|---|
| | Cells EdU positive (%) | Cells EdU negative (%) |
| Control | 0.0 ±0.0 | 100.0 ±0.0 |
| 10 μM EdU | 94.0 ±2.67 | 6.0 ±2.67 |
| 50 μM EdU | 97.67 ±0.44 | 2.33 ±0.44 |
| 100 μM EdU | 97.67 ±1.11 | 2.33 ±1.11 |

| **F.** Day 5 (72 h EdU recovery) | | |
|---|---|---|
| | Cells EdU positive (%) | Cells EdU negative (%) |
| Control | 0.0 ±0.0 | 100.0 ±0.0 |
| 10 μM EdU | 93.67 ±2.44 | 6.33 ±2.44 |
| 50 μM EdU | 95.5 ±2.33 | 4.5 ±2.33 |
| 100 μM EdU | 97.67 ±1.78 | 2.33 ±1.78 |

**Supplementary Table 5.** Numerical sources (mean and standard deviation) of the BrdU/EdU retention time in *Trypanosoma brucei* that were presented in Figure 3C, D.

| **A.** Day 1 (1 h BrdU uptake) | | |
|---|---|---|
| | Cells BrdU positive (%) | Cells BrdU negative (%) |
| Control | 0.0 ±0.0 | 100.0 ±0.0 |
| 10 μM BrdU | 3.33 ±1.11 | 96.67 ±1.11 |
| 50 μM BrdU | 9.17 ±3.44 | 90.83 ±3.44 |
| 100 μM BrdU | 33.17 ±4.56 | 66.83 ±4.56 |

| **B.** Day 2 (24 h BrdU uptake) | | |
|---|---|---|
| | Cells BrdU positive (%) | Cells BrdU negative (%) |
| Control | 0.0 ±0.0 | 100.0 ±0.0 |
| 10 μM BrdU | 7.17 ±1.89 | 92.83 ±1.89 |
| 50 μM BrdU | 12.5 ±3.0 | 87.5 ±3.0 |
| 100 μM BrdU | 39.83 ±11.89 | 60.17 ±11.89 |

| **C.** Day 5 (72 h BrdU recovery) | | |
|---|---|---|
| | Cells BrdU positive (%) | Cells BrdU negative (%) |
| Control | 0.0 ±0.0 | 100.0 ±0.0 |
| 10 μM BrdU | 1.0 ±0.67 | 99.0 ±0.67 |
| 50 μM BrdU | 1.83 ±0.78 | 98.17 ±0.78 |
| 100 μM BrdU | 7.33 ±7.11 | 92.67 ±7.11 |

| **D.** Day 1 (1 h EdU uptake) | | |
|---|---|---|
| | Cells EdU positive (%) | Cells EdU negative (%) |
| Control | 0.0 ±0.0 | 100.0 ±0.0 |
| 10 μM EdU | 16.0 ±4.0 | 84.0 ±4.0 |
| 50 μM EdU | 28.67 ±8.22 | 71.33 ±8.22 |
| 100 μM EdU | 38.5 ±3.67 | 61.5 ±3.67 |

| **E.** Day 2 (24 h EdU uptake) | | |
|---|---|---|
| | Cells EdU positive (%) | Cells EdU negative (%) |
| Control | 0.0 ±0.0 | 100.0 ±0.0 |
| 10 μM EdU | 31.5 ±4.87 | 68.5 ±4.87 |
| 50 μM EdU | 47.47 ±4.31 | 52.53 ±4.31 |
| 100 μM EdU | 83.83 ±2.89 | 16.17 ±2.89 |

| **F.** Day 5 (72 h EdU recovery) | | |
|---|---|---|
| | Cells EdU positive (%) | Cells EdU negative (%) |
| Control | 0.0 ±0.0 | 100.0 ±0.0 |
| 10 μM EdU | 19.5 ±1.33 | 80.5 ±1.33 |
| 50 μM EdU | 34.67 ±4.44 | 65.33 ±4.44 |
| 100 μM EdU | 73.33 ±3.78 | 26.67 ±3.78 |

**Supplementary Table 6.** Numerical sources (mean and standard deviation) of the BrdU/EdU retention time in *Leishmania major* that were presented in Figure 3E, F.

**A.** Day 1 (1 h BrdU uptake)

| | Cells BrdU positive (%) | Cells BrdU negative (%) |
|---|---|---|
| Control | 0.0 ±0.0 | 100.0 ±0.0 |
| 10 µM BrdU | 15.6 ±3.87 | 84.4 ±3.87 |
| 50 µM BrdU | 20.47 ±2.49 | 79.5 ±2.49 |
| 100 µM BrdU | 25.52 ±3.04 | 74.48 ±3.04 |

**B.** Day 2 (24 h BrdU uptake)

| | Cells BrdU positive (%) | Cells BrdU negative (%) |
|---|---|---|
| Control | 0.0 ±0.0 | 100.0 ±0.0 |
| 10 µM BrdU | 49.43 ±5.29 | 50.57 ±5.29 |
| 50 µM BrdU | 82.61 ±6.94 | 17.39 ±6.94 |
| 100 µM BrdU | 87.91 ±3.81 | 12.09 ±3.81 |

**C.** Day 5 (72 h BrdU recovery)

| | Cells BrdU positive (%) | Cells BrdU negative (%) |
|---|---|---|
| Control | 0.0 ±0.0 | 100.0 ±0.0 |
| 10 µM BrdU | 3.45 ±0.56 | 96.55 ±0.56 |
| 50 µM BrdU | 8.2 ±4.27 | 91.8 ±4.27 |
| 100 µM BrdU | 13.7 ±5.73 | 86.3 ±5.73 |

**D.** Day 1 (1 h EdU uptake)

| | Cells EdU positive (%) | Cells EdU negative (%) |
|---|---|---|
| Control | 0.0 ±0.0 | 100.0 ±0.0 |
| 10 µM EdU | 23.33 ±2.44 | 76.67 ±2.44 |
| 50 µM EdU | 38.00 ±2.67 | 62.00 ±2.67 |
| 100 µM EdU | 36.33 ±1.78 | 63.67 ±1.78 |

**E.** Day 2 (24 h EdU uptake)

| | Cells EdU positive (%) | Cells EdU negative (%) |
|---|---|---|
| Control | 0.0 ±0.0 | 100.0 ±0.0 |
| 10 µM EdU | 95.33 ±3.56 | 4.67 ±3.56 |
| 50 µM EdU | 95.33 ±2.22 | 4.67 ±2.22 |
| 100 µM EdU | 97.67 ±1.56 | 2.33 ±1.56 |

**F.** Day 5 (72 h EdU recovery)

| | Cells EdU positive (%) | Cells EdU negative (%) |
|---|---|---|
| Control | 0.0 ±0.0 | 100.0 ±0.0 |
| 10 µM EdU | 68.67 ±5.78 | 31.33 ±5.78 |
| 50 µM EdU | 72.67 ±4.89 | 27.33 ±4.89 |
| 100 µM EdU | 84.0 ±3.33 | 16.0 ±3.33 |

**Supplementary Table 7.** Numerical sources (mean and standard deviation) of the DNA damage signaling detection (using γH2A) in *Trypanosoma cruzi* that were presented in Figure 4A-C.

**A.** Day 1 (1 h BrdU uptake)

| | Cells γH2A positive (%) | Cells γH2A negative (%) | Mean γH2A intensity (au) |
|---|---|---|---|
| Control | 9.33 ±1.56 | 90.67 ±1.56 | 8.21 ±1.95 |
| 10 µM BrdU | 11.33 ±1.11 | 88.67 ±1.11 | 7.78 ±1.82 |
| 50 µM BrdU | 14.0 ±2.67 | 86.0 ±2.67 | 7.44 ±1.84 |
| 100 µM BrdU | 15.0 ±2.0 | 85.0 ±2.0 | 9.92 ±2.08 |

**B.** Day 2 (24 h BrdU uptake)

| | Cells γH2A positive (%) | Cells γH2A negative (%) | Mean γH2A intensity (au) |
|---|---|---|---|
| Control | 9.0 ±1.33 | 91.0 ±1.33 | 6.82 ±1.22 |
| 10 µM BrdU | 12.33 ±1.11 | 87.67 ±1.11 | 7.01 ±2.32 |
| 50 µM BrdU | 13.67 ±1.11 | 86.33 ±1.11 | 6.53 ±1.23 |
| 100 µM BrdU | 15.33 ±0.44 | 84.67 ±0.44 | 6.47 ±1.0 |

**C.** Day 5 (72 h BrdU recovery)

| | Cells γH2A positive (%) | Cells γH2A negative (%) | Mean γH2A intensity (au) |
|---|---|---|---|
| Control | 11.33 ±2.44 | 88.67 ±2.44 | 6.24 ±0.99 |
| 10 µM BrdU | 29.67 ±5.11 | 70.33 ±5.11 | 7.26 ±1.85 |
| 50 µM BrdU | 28.33 ±3.78 | 71.67 ±3.78 | 8.52 ±2.37 |
| 100 µM BrdU | 26.67 ±2.22 | 73.33 ±2.22 | 8.23 ±2.1 |

**D.** Day 1 (1 h EdU uptake)

| | Cells γH2A positive (%) | Cells γH2A negative (%) | Mean γH2A intensity (au) |
|---|---|---|---|
| Control | 8.67 ±1.56 | 91.33 ±1.56 | 6.92 ±1.88 |
| 10 µM EdU | 8.33 ±2.44 | 91.67 ±2.44 | 7.37 ±1.6 |
| 50 µM EdU | 13.00 ±1.33 | 87.0 ±1.33 | 9.11 ±1.99 |
| 100 µM EdU | 10.0 ±2.67 | 90.0 ±2.67 | 10.12 ±1.48 |

**E.** Day 2 (24 h EdU uptake)

| | Cells γH2A positive (%) | Cells γH2A negative (%) | Mean γH2A intensity (au) |
|---|---|---|---|
| Control | 9.0 ±1.33 | 91.0 ±1.33 | 6.21 ±1.7 |
| 10 µM EdU | 14.83 ±0.89 | 85.17 ±0.89 | 11.06 ±1.84 |
| 50 µM EdU | 18.83 ±1.56 | 81.17 ±1.56 | 11.66 ±1.86 |
| 100 µM EdU | 36.67 ±0.89 | 63.33 ±0.89 | 10.65 ±1.5 |

**F.** Day 5 (72 h EdU recovery)

| | Cells γH2A positive (%) | Cells γH2A negative (%) | Mean γH2A intensity (au) |
|---|---|---|---|
| Control | 12.33 ±1.78 | 87.67 ±1.78 | 5.27 ±0.73 |
| 10 µM EdU | 23.67 ±2.89 | 76.33 ±2.89 | 8.9 ±1.8 |
| 50 µM EdU | 72.33 ±3.11 | 27.67 ±3.11 | 13.96 ±3.05 |
| 100 µM EdU | 89.33 ±1.56 | 10.67 ±1.56 | 21.77 ±7.17 |

au: arbitrary units

**Supplementary Table 8.** Numerical sources (mean and standard deviation) of the DNA damage signaling detection (using γH2A) in *Trypanosoma brucei* that were presented in Figure 4D-F.

**A.** Day 1 (1 h BrdU uptake)

| | Cells γH2A positive (%) | Cells γH2A negative (%) | Mean γH2A intensity (au) |
|---|---|---|---|
| Control | 39.67 ±2.78 | 60.33 ±2.78 | 10.03 ±9.4 |
| 10 µM BrdU | 38.50 ±1.33 | 61.50 ±1.33 | 10.58 ±8.36 |
| 50 µM BrdU | 37.83 ±1.56 | 62.17 ±1.56 | 13.9 ±7.39 |
| 100 µM BrdU | 38.67 ±1.89 | 61.33 ±1.89 | 14.72 ±13.7 |

**B.** Day 2 (24 h BrdU uptake)

| | Cells γH2A positive (%) | Cells γH2A negative (%) | Mean γH2A intensity (au) |
|---|---|---|---|
| Control | 37.17 ±3.44 | 62.83 ±3.44 | 10.97 ±7.72 |
| 10 µM BrdU | 38.50 ±1.67 | 61.5 ±1.67 | 10.59 ±7.53 |
| 50 µM BrdU | 40.0 ±3.33 | 60.0 ±3.33 | 14.0 ±9.38 |
| 100 µM BrdU | 42.17 ±2.78 | 57.83 ±2.78 | 21.07 ±15.58 |

**C.** Day 5 (72 h BrdU recovery)

| | Cells γH2A positive (%) | Cells γH2A negative (%) | Mean γH2A intensity (au) |
|---|---|---|---|
| Control | 39.83 ±1.56 | 60.17 ±1.56 | 6.89 ±4.43 |
| 10 µM BrdU | 40.50 ±3.0 | 59.50 ±3.0 | 8.59 ±6.26 |
| 50 µM BrdU | 38.67 ±2.56 | 61.33 ±2.56 | 8.45 ±6.53 |
| 100 µM BrdU | 38.83 ±2.11 | 61.17 ±2.11 | 9.87 ±7.66 |

**D.** Day 1 (1 h EdU uptake)

| | Cells γH2A positive (%) | Cells γH2A negative (%) | Mean γH2A intensity (au) |
|---|---|---|---|
| Control | 38.83 ±2.11 | 61.17 ±2.11 | 11.99 ±8.36 |
| 10 µM EdU | 41.67 ±2.56 | 58.33 ±2.56 | 13.49 ±7.75 |
| 50 µM EdU | 52.0 ±1.67 | 48.0 ±1.67 | 14.89 ±8.62 |
| 100 µM EdU | 51.33 ±5.44 | 48.67 ±5.44 | 18.84 ±14.12 |

**E.** Day 2 (24 h EdU uptake)

| | Cells γH2A positive (%) | Cells γH2A negative (%) | Mean γH2A intensity (au) |
|---|---|---|---|
| Control | 39.83 ±1.78 | 60.17 ±1.78 | 13.19 ±6.34 |
| 10 µM EdU | 39.83 ±3.44 | 60.17 ±3.44 | 10.73 ±6.34 |
| 50 µM EdU | 46.17 ±2.56 | 53.83 ±2.56 | 20.55 ±14.15 |
| 100 µM EdU | 51.33 ±2.78 | 48.67 ±2.78 | 29.56 ±19.41 |

**F.** Day 5 (72 h EdU recovery)

| | Cells γH2A positive (%) | Cells γH2A negative (%) | Mean γH2A intensity (au) |
|---|---|---|---|
| Control | 36.17 ±4.22 | 63.83 ±4.22 | 8.41 ±2.92 |
| 10 µM EdU | 41.17 ±2.56 | 58.83 ±2.56 | 11.35 ±6.74 |
| 50 µM EdU | 40.33 ±1.89 | 59.67 ±1.89 | 11.76 ±4.16 |
| 100 µM EdU | 41.5 ±2.0 | 58.5 ±2.0 | 11.57 ±5.53 |

au: arbitrary units

**Supplementary Table 9.** Numerical sources (mean and standard deviation) of the DNA damage signaling detection (using γH2A) in *Leishmania major* that were presented in Figure 4G-I.

| **A.** Day 1 (1 h BrdU uptake) | | | |
|---|---|---|---|
| | Cells γH2A positive (%) | Cells γH2A negative (%) | Mean γH2A intensity (au) |
| Control | 8.50 ±1.0 | 91.50 ±1.0 | 5.85 ±1.19 |
| 10 μM BrdU | 7.33 ±1.11 | 92.67 ±1.11 | 5.46 ±1.06 |
| 50 μM BrdU | 6.33 ±1.56 | 93.67 ±1.56 | 5.97 ±2.1 |
| 100 μM BrdU | 9.0 ±0.67 | 91.0 ±0.67 | 7.83 ±2.28 |

| **B.** Day 2 (24 h BrdU uptake) | | | |
|---|---|---|---|
| | Cells γH2A positive (%) | Cells γH2A negative (%) | Mean γH2A intensity (au) |
| Control | 5.33 ±1.56 | 94.67 ±1.56 | 5.79 ±0.94 |
| 10 μM BrdU | 8.83 ±1.89 | 91.17 ±1.89 | 6.75 ±1.8 |
| 50 μM BrdU | 8.33 ±1.56 | 91.67 ±1.56 | 6.27 ±1.19 |
| 100 μM BrdU | 6.33 ±1.11 | 93.67 ±1.11 | 8.03 ±1.41 |

| **C.** Day 5 (72 h BrdU recovery) | | | |
|---|---|---|---|
| | Cells γH2A positive (%) | Cells γH2A negative (%) | Mean γH2A intensity (au) |
| Control | 8.0 ±0.67 | 92.0 ±0.67 | 8.09 ±1.71 |
| 10 μM BrdU | 7.83 ±1.44 | 92.17 ±1.44 | 8.01 ±1.88 |
| 50 μM BrdU | 9.67 ±1.78 | 90.33 ±1.78 | 9.26 ±2.53 |
| 100 μM BrdU | 14.67 ±3.11 | 85.33 ±3.11 | 9.58 ±2.83 |

| **D.** Day 1 (1 h EdU uptake) | | | |
|---|---|---|---|
| | Cells γH2A positive (%) | Cells γH2A negative (%) | Mean γH2A intensity (au) |
| Control | 12.0 ±1.33 | 86.67 ±1.33 | 6.08 ±1.86 |
| 10 μM EdU | 19.0 ±2.67 | 81.0 ±2.67 | 6.47 ±2.24 |
| 50 μM EdU | 19.33 ±1.11 | 80.67 ±1.11 | 8.11 ±2.79 |
| 100 μM EdU | 17.33 ±2.22 | 82.67 ±2.22 | 11.45 ±2.94 |

| **E.** Day 2 (24 h EdU uptake) | | | |
|---|---|---|---|
| | Cells γH2A positive (%) | Cells γH2A negative (%) | Mean γH2A intensity (au) |
| Control | 10.0 ±0.67 | 90.0 ±0.67 | 6.93 ±2.13 |
| 10 μM EdU | 61.0 ± 5.33 | 39.0 ± 5.33 | 13.32 ±6.73 |
| 50 μM EdU | 64.0 ±2.67 | 36.0 ±2.67 | 17.14 ±7.29 |
| 100 μM EdU | 61.67 ±2.22 | 38.33 ±2.22 | 16.52 ±7.58 |

| **F.** Day 5 (72 h EdU recovery) | | | |
|---|---|---|---|
| | Cells γH2A positive (%) | Cells γH2A negative (%) | Mean γH2A intensity (au) |
| Control | 9.0 ±1.33 | 91.0 ±1.33 | 5.88 ±1.72 |
| 10 μM EdU | 34.0 ±4.0 | 66.0 ±4.0 | 7.68 ±5.07 |
| 50 μM EdU | 31.0 ±3.33 | 69.0 ±3.33 | 7.45 ±2.33 |
| 100 μM EdU | 38.33 ±2.89 | 61.67 ±2.89 | 8.0 ±3.42 |

au: arbitrary units
